# The metagenomic analysis of viral diversity in Colorado potato beetle public NGS data

**DOI:** 10.1101/2023.01.04.522816

**Authors:** Maria Starchevskaya, Ekaterina Kamanova, Yuri Vyatkin, Tatyana Tregubchak, Tatyana Bauer, Sergei Bodnev, Ulyana Rotskaya, Olga Polenogova, Vadim Kryukov, Denis Antonets

## Abstract

The Colorado potato beetle (CPB) is one of the most serious insect pests with high ecological plasticity and ability to rapidly develop resistance to insecticides. The use of biological insecticides based on viruses is a promising approach to control insect pests, but the information on viruses, which infect leaf feeding beetles, is scarce. We performed the metagenomic analysis of 297 CPB genomic and transcriptomic samples from public NBCI SRA database. The reads that were not aligned to the reference genome were assembled with metaSPAdes and 13314 selected contigs were analyzed with BLAST tools. The contigs and non-aligned reads were also analyzed with Kraken2 software. 3137 virus-positive contigs were attributed to different viruses belonging to 6 types, 17 orders and 32 families, matching over 97 viral species. The annotated sequences can be divided into several groups: homologous to genetic sequences of insect viruses (*Adintoviridae, Ascoviridae, Baculoviridae, Dicistroviridae, Chuviridae, Hytrosaviridae, Iflaviridae, Iridoviridae, Nimaviridae, Nudiviridae, Phasmaviridae, Picornaviridae, Polydnaviriformidae, Xinmoviridae* etc.), plant viruses (*Betaflexiviridae, Bromoviridae, Kitaviridae, Potyviridae*), and endogenous retroviral elements (*Retroviridae, Metaviridae*). Also, the full-length genomes and near-full length genome sequences of several viruses were assembled. We have also found the sequences belonging to Bracoviriform viruses and for the first time experimentally validated the presence of bracoviral genetic fragments in CPB genome. Our work is the first attempt to discover the viral genetic material in CPB samples and we hope that further studies will help to identify new viruses to extend the arsenal of biopesticides against CPB. The analytical pipeline and additional materials are available at https://github.com/starchevskaya-maria17/uncoVir

## 1. Introduction

The metagenomic approach has proven to be a highly efficient way of detecting viruses in a variety of environments. Unlike traditional cultural and diagnostic methods, viral metagenomics uses high-throughput next-generation sequencing, which makes it possible to identify rare and new viruses and to establish ecological links between microorganisms and their natural habitat [1]. Among the various routes of virus transmission, insect transmission is one of the most widespread. Insects are the largest group of animals on the planet and are of significant ecological, agricultural, and medical importance. However, our knowledge of insect viruses is still limited. Establishing an insect virome will provide new insights into the ecology and evolution of viruses and help address public health issues in the fight against arbovirus infections. In addition, pathogenic insect viruses can be used for the biological control of insect pests. Thus, identifying new insect viruses can expand the arsenal of potential biocontrol agents [2].

The Colorado potato beetle is one of the most widespread and destructive potato pests in the world. Its ability to adapt to various *Solanaceae* plants, high ecological plasticity, flexible life-cycle, and rapid development of insecticide resistance have led to its global expansion [3]. In addition, leaf-feeding beetles transmit a variety of plant viruses. *Leptinotarsa decemlineata* was shown to transmit the Potato virus Y, which causes the necrosis of leaves and tubers and can significantly reduce the yield and storage time of potato crops. Moreover, the Potato virus Y reduces the production of potato defense factors, such as sesquiterpenes, namely beta-barbatene, thus increasing the growth of Colorado potato beetle larvae feeding on infected plants. As a result, the plants are exposed to a combination of severe biotic stressors, increasing crop losses [4]. During its lifetime, the Colorado potato beetle female can lay up to 800 eggs and produce up to three generations per year, depending on climatic conditions [5]. Unless insecticides are used, the Colorado potato beetle can cause 40–80% of potato crop losses. Although the use of insecticides has led to a sharp decline in populations, the development of resistance to all chemicals registered to date has been observed. Increasing the dosage provides a short-term improvement while significantly increasing the rate of resistance development [6]. Several research groups have established the reasons for the rapid evolutionary changes when studying the genetic basis of the resistance of the Colorado potato beetle to insecticides and its adaptation to environmental factors [3]. Given that approximately 17% of its genome is composed of mobile genetic elements and that there is a high level of nucleotide diversity in fast-growing populations, the Colorado potato beetle evolves faster than many other beetles [7]. Its high ability to acclimatize is related to specific genes associated with the switch of metabolism, growth, and diapause of insects [3]. The mechanisms of detoxification of secondary plant compounds and, probably, insecticidal compounds are associated with a wide range of highly variable genes, including genes for carboxylesterase, glutathione-S-transferase, and cytochrome P450, with the adaptation to feeding on various *Solanaceae* plants involving the expansion of genes encoding the digestive enzymes and bitter taste receptors [7]. Colorado potato beetles are expected to develop resistance to all newly introduced insecticides [6]. In addition, the widespread use of chemical pesticides can lead to serious environmental problems because of their nonspecific effects on many other animal species.

Biological control methods are the most environmentally friendly ways to eliminate various pests of agricultural crops. The release of biological control agents to manage pest populations has great potential, but only a few natural enemies could be massively grown at scale. In addition, the development of a natural enemy population is often slower than that of the target insect, usually requiring different conditions for optimal growth [8]. Currently, the existing biological control methods use pathogenic fungi, such as *Beauveria bassiana, Metarhizium* species [9] and bacteria *Bacillus thuringiensis* [10], with no biological control agents based on viruses. The only exceptions are the preparations based on viruses of the family *Baculoviridae*, which are widely used to control insect pests of the order *Lepidoptera* [11].

Currently, over 20 virus families are known to affect different groups of insects. Most insect virus families cause arbovirus infections and, together with plant viruses, can be transmitted to humans [12]. Nevertheless, many insect viruses are not dangerous to humans. They are used for controlling insect pests, as model objects and as platforms for developing vector systems [2].

Leaf beetles of the family *Chrysomelidae* are assumed to be affected by viruses of the families *Baculoviridae* and *Iridoviridae* [13]. And the literature sources on viruses affecting *L. decemlineata* is extremely scarce. Despite being unquestionably significant for agriculture, the Colorado potato beetle virome is practically unstudied and our study is the first attempt to detect viral genetic material in the genomic and transcriptomic data of *L. decemlineata* samples (obtained from the NCBI SRA database). Such studies could help to improve the understanding of the Colorado potato beetle ecology and may help to identify new viruses to enrich the arsenal of biopesticides and eventually reduce the economic losses in agriculture.

## 2. Materials and Methods

### Genome and transcriptome data of *Leptinotarsa decemlineata* samples

The data on 307 samples of *L. decemlineata* were obtained from the NBCI SRA database, with 297 samples selected for the study, including the genomic data obtained from muscle tissues (PRJNA508767, PRJNA369863, PRJNA580490) and whole insect tissues (PRJNA171749); the transcriptomic data obtained from antennae tissues (PRJNA280017), heads (PRJNA400685), intestines (PRJNA400685, PRJNA336167), larvae (PRJNA275431, PRJNA694179), and whole insect tissues (PRJNA438159, PRJNA384383, PRJNA297027, PRJNA464380, PRJNA275662, PRJNA353242, PRJNA646009, PRJNA553565). The sample sizes of the transcriptomic data varied from 217 Mbp to 20.5 Gbp, with genomic data ranging from 7.1 Gbp to 20.5 Gbp. The genomic samples were sequenced with Illumina HiSeq 2000 (PRJNA508767, PRJNA369863, and PRJNA171749) and Illumina HiSeq 1500 (PRJNA580490). The transcriptomic samples were sequenced with Illumina HiSeq 4000 (PRJNA438159), Illumina HiSeq 2000 (PRJNA280017, PRJNA400685, PRJNA400685, and PRJNA336167), Illumina HiSeq 2500 (PRJNA275431, PRJNA438159, PRJNA384383, PRJNA297027, PRJNA464380, PRJNA275662, PRJNA353242, and PRJNA646009), and Illumina NextSeq 500 (PRJNA553565). The primary data sources are presented in Supplementary Table 1. Project PRJNA171749 [7] was aimed at assembling and annotating the *L. decemlineata* genome, and the data from the PRJNA400685 and PRJNA297027 [14] projects was primarily used to assemble and to annotate the Colorado potato beetle transcriptome.

### Databases

1. The NCBI SRA database (https://www.ncbi.nlm.nih.gov/sra).
2. The BigViralDB – our custom database – combined a nonredundant viral sequences assembled from four databases: NCBI RefSeq (11566 viral sequences, June 2021) (https://www.ncbi.nlm.nih.gov/refseq/), Virxicon (327932 viral sequences, June 2021) [15], Virosaurus (823421 viral sequences, April 2020) [16], and virus sequences from the NCBI Nucleotide database (1367485 viral sequences, June 2021). The database could be provided at request.
3. The NCBI Genome Database (https://www.ncbi.nlm.nih.gov/genome/, July 2020).
4. The NCBI nt database (https://ftp.ncbi.nlm.nih.gov/blast/db/FASTA/nt.gz, July 2020).
5. The NCBI nr database (https://ftp.ncbi.nlm.nih.gov/blast/db/FASTA/nr.gz, July 2020).
6. The NCBI UniVec database (https://www.ncbi.nlm.nih.gov/tools/vecscreen/univec/), a collection of unique vector sequences used in genetic engineering and biotechnology research (3137 sequences, June 2021).

### Metagenome assembly and sequence classification

We constructed an analytical pipeline to analyze the genomic and transcriptomic data of *Leptinotarsa decemlineata* obtained from the NCBI SRA database (Fig. 1). The pipeline, written in Snakemake [17], involves several approaches for classifying genetic sequences and is freely available at: https://github.com/starchevskayamaria17/uncoVir.

**Figure 1.**
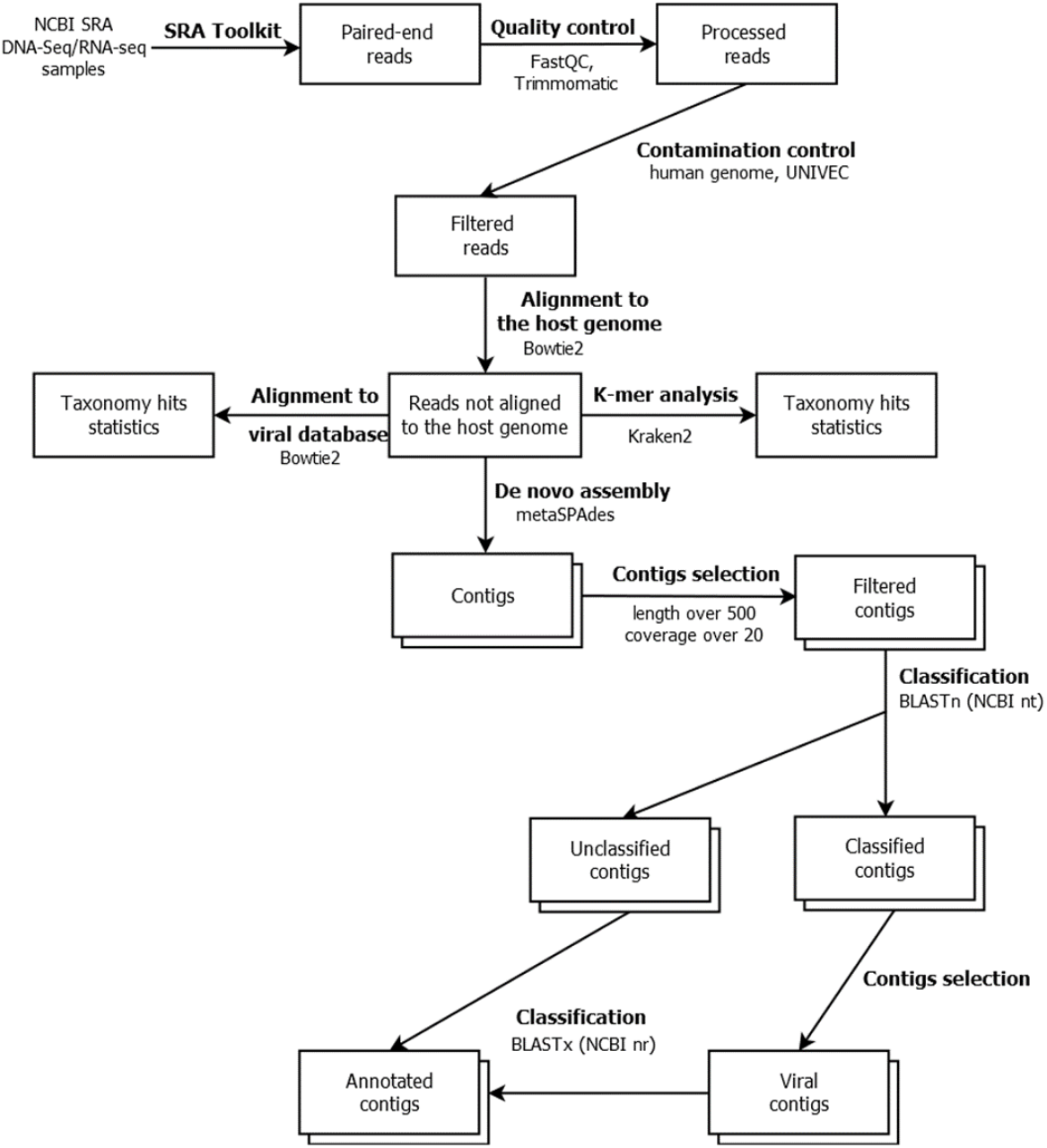
The software pipeline used in this work, with the analysis steps and their connections indicated and the programs used specified. https://github.com/starchevskayamaria17/uncoVir.

In the first step, low-quality and short reads (quality less than 20 and length less than 30 nucleotides), overrepresented sequences, and adapters were removed using the Trimmomatic tool [18]. For contamination control, the reads aligned to the human genome (GRCh38.p12) using the Bowtie2 tool were deleted [19]. The reads of synthetic origin (identified with BLAST against the UniVec database) were also deleted. The reads aligned to the reference *L. decemlineata* genome (assembly GCA_000500325.2 Ldec_2.0) using the Bowtie2 tool were also excluded from the subsequent analysis.

In the next step, viral diversity was assessed using the k-mer spectrum analysis. The unaligned reads were classified with Kraken2 [20] using our customly compiled viral sequence database BigViralDB. The results were visualized using Pavian [21]. To assess the completeness of particular viral genomes, we have also aligned the reads using Bowtie2 to the viral genomic sequences contained in our BigViralDB database.

The reads unaligned on the *L. decemlineata* genome were assembled into contigs using the SPAdes assembler tool [22]. Contigs longer than 500 nucleotides and with coverage of more than 20 were selected and analyzed using BLASTn against the NBCI Nucleotide database. The unclassified contigs and contigs annotated with BLASTn as virus-positive were then analyzed using BLASTx against the NCBI nonredundant protein database. The obtained annotation results were analyzed and visualized using custom scripts written in Python programming language. The details of the running parameters of the programs were included in the pipelines and are given in the source code in the repository (https://github.com/starchevskayamaria17/uncoVir).

### Analysis of reference genomes of selected coleopterans

The genome sequence of *Leptinotarsa decemlineata* (assembly GCA_000500325.2 Ldec_2.0) used as a reference for alignment was split into 31-nucleotide long k-mers and analyzed using the BLASTn tool against the NBCI Nucleotide database. Similarly, other reference genomes of Coleoptera obtained from the NCBI Genome database were examined: *Diabrotica virgifera* (GCA_003013835.2 Dvir_v2. 0), *Oryzaephilus surinamensis* (GCA_004796505.1 ASM479650v1), *Coccinella septempunctata* (GCA_003568925.1 Csep_MD8_v1), *Harmonia axyridis* (GCA_011033045. 1), *Aleochara bilineata* (GCA_003054995.1 ASM305499v1), *Aethina tumida* (GCA_001937115.1 Atum_1.0), *Oryctes borbonicus* (GCA_902654985. 1), *Nicrophorus vespilloides* (GCA_001412225.1 Nicve_v1.0), *Asbolus verrucosus* (GCA_004193795.1 BDFB_1.0), *Protaetia brevitarsis* (GCA_004143645. 1 ASM414364v1), *Popillia japonica* (GCA_004785975.1 GSC_JBeet_1), *Anoplophora glabripennis* (GCA_000390285.2 Agla_2. 0), *Pogonus chalceus* (GCA_002278615.1 Pchal_1.0), *Agrilus planipennis* (GCA_000699045.2 Apla_2.0), *Sitophilus oryzae* (GCA_002938485.2 Soryzae_2. 0), *Onthophagus taurus* (GCA_000648695.2 Otau_2.0), *Dendroctonus ponderosae* (GCA_000355655.1), *Tribolium castaneum* (GCA_000002335.3 Tcas5.2).

### Biological samples collection and preparation

The experiments were carried out on biological samples (*L. decemlineata* adults, larvae, and eggs) collected from private potato fields free from treatment by biological insecticides (Novosibirsk region, Russian Federation; 53°44’3.534”N, 77°39’0.0576”E). With these potato fields not located in protected areas, there was no need for special permission to collect beetles. Landowners did not prevent access to the fields. Endangered or protected species were not used in this work. The insects were kept in a ventilated laboratory room for a 12:12 h dark/light period at a constant temperature of 25 °C. The insects were fed with fresh *Solanum tuberosum* foliage. Fourth-instar larvae (2–4 h postmolt in IV instar) were used for cuticle and hemolymph sample collection. The cuticle was cleared of the fat body in a cold phosphate buffer and frozen in liquid nitrogen, and each sample contained the cuticle of 6 insects. Hemolymph samples were collected as follows. A puncture was made with a sterile needle, and 25 μl of hemolymph was taken. The samples were cooled on ice and frozen in liquid nitrogen, each taken from a single insect [23]. For a more detailed description of biological sample collection and preparation, one is referred to the following papers [24,25].

Colorado potato beetle egg samples were prepared as follows. Ten eggs from three clutches were pulled in a single 1.5 ml Eppendorf tube (1 sample contained 30 eggs). Then, 0.5 ml of 6% hydrogen peroxide was added to the sample, the sample was stirred for 5 minutes at 500 rpm on a TS-100C (BioSan) shaker, and the free liquid was removed. Then, 0.5 ml of autoclaved water was added to the sample, the tube was inverted several times, and the remaining liquid was removed with a sterile filter paper. Then, the procedure was repeated with 2.5% sodium hypochlorite. All manipulations were performed at room temperature with room temperature reagents, with the shaker used without heating. The prepared samples were frozen in liquid nitrogen. This method was adapted from the paper [23].

### Confirmation of bracovirus genetic fragments presence in the genetic material of *L. decemlineata*

For the two fragments classified as bracovirus fragments, the oligonucleotide primers were calculated using Primer3 [26]. The sequences of the fragments and primers are presented in the supplementary materials in Supplementary Table 2.

DNA was extracted from 20 mg of sterile eggs, cuticle, hemolymph, and whole insect (imago) of the Colorado potato beetle using the PureLink genomic DNA mini kit (Invitrogen, United States). The amplification reaction was performed in a GeneAmp PCR System 9700 thermal cycler (Applied Biosystems, USA) in a volume of 50 μl. The reaction mixture contained Taq-DNA polymerase buffer (Invitrogen, USA), 1.5 mM MgCl2, 0.2 mM dTTP, 0.2 mM dGTP, 0.2 mM dATP, 0.2 mM dCTP, 10 pmol of each oligonucleotide primer, 1.25 Taq-DNA polymerase activity units (Invitrogen, USA), and 2–10 ng matrix DNA. The amplification was performed for 30 cycles using a stepwise program (94°, 45 sec; 55°, 45 sec; 72°, 2 min). The PCR products were purified using a PCR Purification Kit (250) (Qiagen, Germany). The amplification products were separated in a 1.2 % agarose gel. The fragments obtained were confirmed using Sanger sequencing. The reaction was performed in a GeneAmp PCR System 9700 thermal cycler (Applied Biosystems, USA) in a volume of 5 μl. The reaction mixture contained prepurified DNA matrix, 2 μl of BigDye-v.3.1 sequencing solution (PE BioSystems, USA), and a 4 pM oligonucleotide primer for 30 cycles using a stepwise program (94°, 20 sec; 50°, 20 sec; 60°, 4 min).

## 3. Results

### Analysis of the k-mer spectra

The genomic and transcriptomic data of 297 *L. decemlineata* samples was analyzed in this study. The quality and contamination control of the reads for each sample resulted in no more than 5% of the reads being removed. Up to 10% of the reads were not aligned to the *L. decemlineata* genome. Among the reads not aligned to the *L. decemlineata* genome, 1.36% of the reads from the genomic samples (8,461,508 of 620,747,359) and 3.04% of the reads from the transcriptomic samples (5,905,253 of 193,959,828) were classified using Kraken2 and our custom BigViralDB database. The results of the analysis down to the family level are visualized in Figure 2. In the transcriptomic samples, the viral reads were classified as mainly belonging to the viral families of *Betaflexviridae, Baculoviridae, Peribunyaviridae, Potyviridae, Orthomyxoviridae, Paramyxoviridae, Dicistroviridae, Polydnaviridae*, and others. In contrast to the transcriptomic data, the genomic data lacked the reads attributed to viral families of *Betaflexviridae, Potyviridae*, and *Dicistroviridae*. The sequences attributed as retroviral or bacteriophage-related are not represented in the plots.

**Figure 2.**
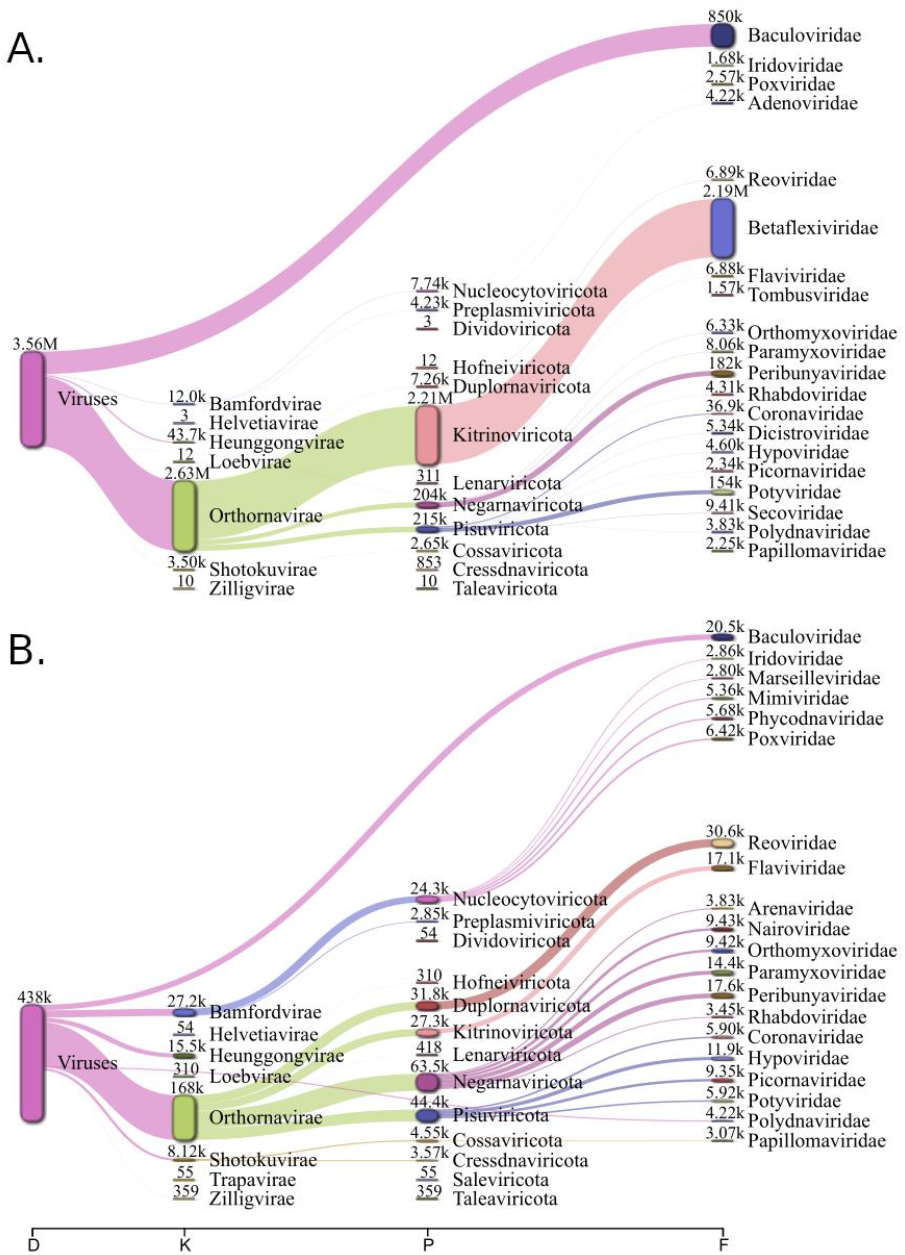
Results of k-mer spectra analysis. The classification of reads attributed as viral was performed down to the family level. The aggregated transcriptomic data analysis is shown on the top chart (A) and the genomic data analysis – on the bottom (B). The numbers of reads are shown matched at each level.

### Contigs analysis

A total of 31,069,568 contigs were collected from all samples as a result of the assembly of reads unaligned to the *L. decemlineata* genome. Contig filtering by length and coverage resulted in the selection of 122,506 contigs (with the average length of 1,861 bp and the maximum contig length of 618,970 bp), which were further classified using BLAST tools. The final table (Supplementary Table 3) with all the results of contigs annotation with BLASTx contains 5,175,489 lines (the e-value < 10^-3^ was chosen as the filtering criterion; matches with bacteriophage proteins were excluded from the analysis). This table provides information on 13,314 unique contigs: the superkingdom Eukaryota accounts for 13,113 unique contigs, the superkingdom Bacteria accounts for 6,220, Archaea accounts for 269, and viruses account for 3,137 contigs. A total of 719 contigs annotated with BLASTn as viral were then annotated with BLASTx. Among the contigs that could not be annotated with BLASTn, 3,395 contigs were assembled from Colorado potato beetle transcriptomic reads and 9,203 – from genomic reads and then they were annotated with BLASTx.

The following tables were created based on the results of BLASTx annotation:

- Among the viral protein BLASTx hits, for which the virus family was known, the best match was selected for each contig based on the e-value (minimum) and the bitscore (maximum), – 2,954 rows (Supplementary Table 4);
- Among the viral protein BLASTx hits, for which the virus family was unknown, the best match was selected for each contig based on the e-value (minimum) and the bitscore (maximum) – 1,662 rows (Supplementary Table 5);
- Among the non-viral protein BLASTx hits the best match was selected for each contig based on the e-value (minimum) and the bitscore (maximum) – 13,252 rows (Supplementary Table 6).

Supplementary Table 7 presents the results of the virus-positive contig annotation after filtering: 1,652 contigs assigned to different virus species with an unspecified virus family, and 2,954 contigs with an established virus family (with the information on contigs the protein products of which have homology to bacteriophage proteins being excluded). This table also provides the identifiers of genomic and transcriptome data projects, the information on protein products, the distribution of contig and alignment lengths, identity scores, and e-value values. Table 1 presents the most interesting results. Virus-positive contigs belong to 6 virus types: *Artverviricota, Kitrinoviricota, Negarnaviricota, Nucleocytoviricota, Pisuviricota, Preplasmiviricota*, distributed in 17 orders, 32 viral families, and 97 species. In addition, at least 48 virus species with unknown viral family were also matched. Figure 3 shows the distributions of virus-positive contigs numbers derived from genomic and transcriptomic samples by virus family or species (if family is unknown). If homology with viral sequences was not taken into account, the most of these contigs were attributed to insects (Coleoptera) as it is shown in Supplementary figures SF1-2.

**Figure 3.**
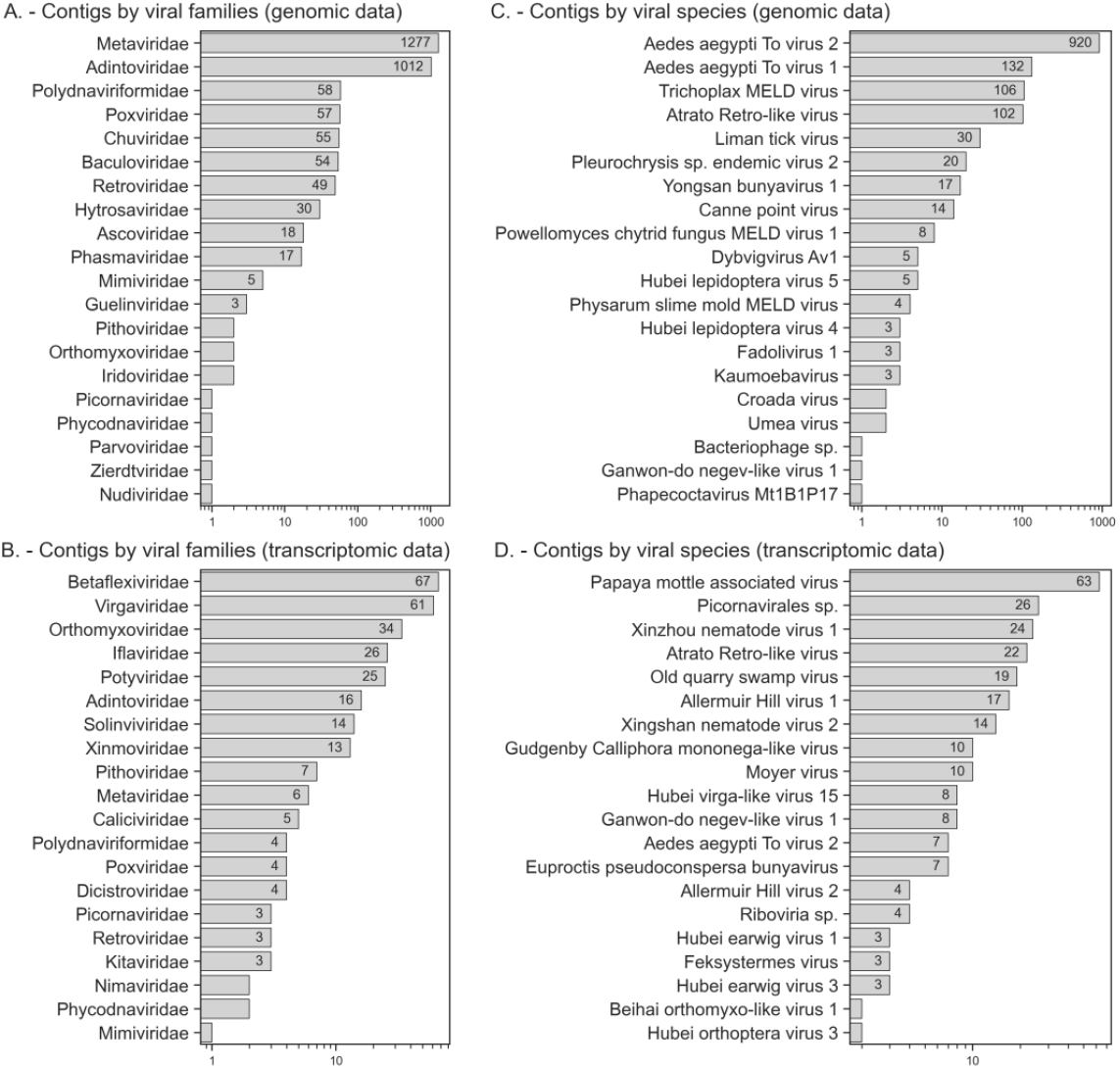
The number of virus-positive contigs by virus family in genomic (A) and transcriptomic (B) samples. The distribution of virus-positive contigs by virus species (unknown family) in the genomic (C) and transcriptomic (D) samples.

**Table 1.** The number of contigs by viral species, family and order, with maximum alignment length and alignment identity percentage, and maximum contig length. The complete table is given in Supplementary Table 7 with project identifiers, protein products, extended taxonomy, alignment e-values etc.

| Order | Family | Species | Max alignment length, aa | Max identity, % | Max contig length, bp | Number of contigs | Protein product |
| --- | --- | --- | --- | --- | --- | --- | --- |
| Martellivirales | Virgaviridae | Nephila clavipes virus 4 | 1047 | 40.51 | 10559 | 51 | RdRp |
|  |  | Broome virga-like virus 1 | 177 | 31.64 | 2499 | 5 | ORF3 |
|  |  | Insect virga-like virus 1 | 165 | 41.67 | 540 | 2 | Hypothetical protein |
| Tymovirales | Betaflexiviridae | Potato virus S | 1975 | 100 | 8539 | 66 | Structural and nonstructural polyprotein |
| Bunyavirales | Phasmaviridae | Wuhan mosquito orthophasmavirus 1 | 388 | 27.06 | 1667 | 17 | Glycoprotein |
| Articulavirales | Orthomyxoviridae | Quaranjavirus araguariense | 437 | 27.82 | 1718 | 9 | Hemagglutinin |
|  |  | Photinus pyralis orthomyxo-like virus 1 | 768 | 43.94 | 2529 | 15 | Nucleocapsid protein, polymerase PB2 |
|  |  | Coleopteran orthomyxo-related virus OKIAV184 | 501 | 31.34 | 1715 | 2 | Hemagglutinin |
|  |  | Coleopteran orthomyxo-related virus OKIAV186 | 761 | 46.78 | 2607 | 2 | Polymerase PB2 |
| Jingchuvirales | Chuviridae | Nigecruvirus ixodes | 631 | 36.42 | 2176 | 14 | Glycoprotein |
|  |  | Coleopteran chu-related virus OKIAV127 | 167 | 72.79 | 1926 | 22 |  |
|  |  | Orthopteran chu-related virus OKIAV152 | 138 | 38.41 | 2075 | 1 |  |
| Mononegavirales | Xinmoviridae | Orthopteran anphevirus | 383 | 25.10 | 1645 | 3 | Hypothetical protein 1 |
|  |  | Ulegvirus freckenfeldense | 1930 | 30.95 | 14588 | 9 | RdRp |
| Pimascovirales | Ascoviridae | Trichoplusia ni ascovirus 2c | 137 | 40.63 | 1321 | 19 | Hypothetical protein |
| Pimascovirales | Iridoviridae | Iridovirus LCIVAC01 | 168 | 30.95 | 796 | 2 | Uncharacterized protein |
| Picornavirales | Caliciviridae | Soybean thrips Pernambuco virus | 497 | 25.81 | 11259 | 6 | Polyprotein |
|  | Dicistroviridae | Aphid lethal paralysis virus | 808 | 92.82 | 3468 | 1 | Capsid protein |
|  |  | Aphis gossypii virus | 1039 | 93.94 | 6635 | 1 | ORF1 |
|  |  | Bivalve RNA virus G5 | 369 | 24.93 | 6436 | 1 | Replicative protein |
|  | Iflaviridae | Lampyris noctiluca iflavirus 1 | 1642 | 39.73 | 9934 | 26 | Putative polyprotein |
|  | Soliniviridae | Solenopsis invicta virus 3 | 966 | 29.12 | 11648 | 12 | Structural and nonstructural polyprotein |
| Patatavirales | Potyviridae | Potato virus Y | 3061 | 100 | 9728 | 25 | Coat protein, polyprotein |
| Lefavirales | Baculoviridae | Mythimna unipuncta nucleopolyhedrovirus | 309 | 25.21 | 1929 | 14 | F-protein |
|  |  | Peridroma alphabaculovirus | 474 | 30.95 | 2668 | 9 |  |
|  |  | Mythimna unipuncta granulovirus B | 251 | 25.10 | 1628 | 14 |  |
|  |  | Helicoverpa armigera nucleopolyhedrovirus | 276 | 22.60 | 1550 | 4 | Envelope protein, hypothetical protein |
|  |  | Lambdina fiscellaria nucleopolyhedrovirus | 156 | 35.90 | 962 | 1 | Vp80 |
|  |  | Plutella xylostella granulovirus | 271 | 21.03 | 1275 | 1 | PxORF26 peptide |
|  |  | Xestia c-nigrum granulovirus | 220 | 26.82 | 1394 | 1 | ORF27 |
|  | Nudiviridae | Penaeus monodon nudivirus | 655 | 28.55 | 17638 | 1 | P74 |
|  | Polydnaviriformidae | Bracoviriform congregatae | 242 | 71.43 | 1669 | 14 | Structural and nonstructural protein |
|  |  | Bracoviriform inaniti | 419 | 43.46 | 2975 | 28 |  |
|  |  | Cotesia sesamiae bracovirus | 614 | 54.76 | 6071 | 19 |  |
|  |  | Bracoviriform facetosae | 158 | 25.95 | 716 | 1 |  |
|  |  | Wuhan insect virus 33 | 1039 | 99.32 | 6635 | 1 | Hypothetical protein |
|  |  | Papaya mottle associated virus | 1017 | 64.40 | 8539 | 63 | RdRp |
|  |  | Old quarry swamp virus | 794 | 46.36 | 2607 | 20 | Putative nucleocapsid, polymerase |
|  |  | Allermuir Hill virus 1 | 1016 | 46.63 | 9840 | 17 | Putative non-structural polyprotein |

The highest number of annotated virus-positive contigs belonged to the endogenous viral elements of retroviral origin: species of *Aedes aegypti To virus 2* (*AAToV2*), *Aedes aegypti To virus 1* (*AAToV1*) and *Atrato Retro-like virus* with unspecified taxonomic affiliation and a number of virus representatives of the families *Metaviridae* and *Retroviridae*. The identity percentage ranges from 31.5% to 85.7% and is higher for the members of the *Metaviridae family, AAToV1* and *AAToV2*. For *Trichoplusia ni TED virus (Metaviridae*), the maximum alignment length reaches 1157 aa. More than 1000 contigs obtained mainly from genomic data were annotated as belonging to different members of the family *Adintoviridae*. The alignments indicate the homology not only to retrovirus-like integrase and polymerase B but also to structural proteins of the family *Adintoviridae*.

Many virus-positive contigs were found to be associated with viruses infecting insects. Twenty-six contigs with similarity to *Lampyris noctiluca iflavirus 1* (family *Iflaviridae*) were identified. The maximum length of alignments between the proteins encoded in these contigs and the polyprotein of this virus was 1642 aa (median 1638 aa), with a median identity of 33.8% (maximum identity percentage 39.7%). The genome size of the *Iflaviridae* family members varies from 9 to 11 kb, with a median length among the corresponding virus-positive contigs being 9870 nucleotides (maximum 9934). Two of the four contigs assigned to the family *Dicistroviridae* were attributed to the species *Aphis gossypii virus* and *Aphid lethal paralysis virus*, with the contig lengths being of 6635 and 3468 nucleotides, respectively, and the genome sizes of representatives of the family *Dicistroviridae* ranging from 8 to 10 kb. The percentage of alignment identity predicted for these contigs is 93.9% for the ORF1 protein (alignment length 1039 aa) and 92.8% for the capsid protein (alignment length 808 aa). Along with the families *Iflaviridae* and *Dicistrovirida*e, other virus-positive contigs homologous to the genetic sequences of the order *Picornavirales* were assigned to insect-infecting members of the *Picornaviridae, Solinviviridae*, and *Caliciviridae* families. The identity percentage of the proteins encoded by these contigs versus the polyproteins of these virus families does not exceed 30%. Many of these contigs are extended and range from 3350 to 11648 nucleotides in length. The protein products of 36 contigs annotated as belonging to insect-infecting members of the family *Orthomyxoviridae* (*Photinus pyralis orthomyxo-like virus 1, Quaranjavirus araguariense, Coleopteran orthomyxo-related viruses*, etc.) showed homology with various structural and nonstructural proteins, with identities of 25 to 55% and an alignment length reaching 768 aa.

55 contigs derived from genomic samples were annotated as belonging to the family *Baculoviridae*, with most contigs encoding polypeptides homologous to F-protein (with a maximal alignment length of 474 aa and maximal identity of 30.9%). In addition, translation frames were identified, with their products homologous to vp80 (156 aa alignment length, 35.9% identity), ORF27 (220 aa alignment length, 26.8% identity), and PxORF26 (271 aa alignment length, 21% identity). Additionally, more than 70 contigs containing translation frames homologous to glycoproteins from various members of the families *Chuviridae* and *Phasmaviridae* were also detected. The identity of the encoded amino acid sequences to the Coleopteran chu-related virus glycoprotein OKIAV127 (family *Chuviridae*) was as high as 72.8%. A total of 62 contigs up to 6 kb in length were identified, with their predicted proteins being homologous with various amino acid sequences of representatives of the genus *Bracoviriform* (family *Polydnaviriformidae*). Additionally, a contig with a length of 17638 nucleotides encoding a protein identical by 28.6% to the protein P74 of *Penaeus monodon nudivirus* (family *Nudiviridae*) was found with an alignment length of 655 aa.

Additionally, we also identified the contigs attributed to plant viruses. Within 66 extended contigs exceeding 8 kb in length, translation frames encoding proteins with 100% identity to *Potato virus S* (family *Betaflexiviridae*) were found, with an alignment length of 1975 aa. A contig was found with a protein product 99.6% homologous to the *Potato virus H* (family *Betaflexiviridae*) shell protein, with an alignment length of 292 aa. Twenty-five contigs longer than 9 kb were found to be assigned to *Potato virus Y* (family *Potyviridae*), with an alignment length of 3061 aa (polyprotein). Given that the genomes of these potato viruses range from 5.4 kb to 12 kb in length, we have obtained almost complete genome assemblies. In addition, we identified 63 extended (up to 8.5 kb) contigs belonging to the *Papaya mottle associated virus*, with their protein products being homologous to RdRp of the virus specified (maximal identity was 64.4%, maximal alignment length was 1017 aa). Additionally, we identified single contigs of plant viruses with protein products being homologous to those of *Garlic common virus*, *Bromoviridae*, and *Kitaviridae* (with identity less than 50%, and the alignment length less than 200 aa).

In addition, contigs were found encoding protein products homologous to viral proteins of the replication system, such as RdRp, and to uncharacterized viral proteins, with identity of 30-40% and alignment length not exceeding 300 aa. In particular, 19 contigs identified in the study were attributed to *Trichoplusia ni ascovirus 2c* (family *Ascoviridae*), the alignment identity of the encoded protein products to various uncharacterized viral proteins was 40.6%, with a maximum alignment length of 137 aa. In the genomic samples we have also identified 2 contigs, whose protein products were 31% identical to the short amino acid sequences of 168 aa in length of the uncharacterized *Iridovirus LCIVAC01* proteins (family *Iridoviridae*). Extended contigs of more than 10 kb in length were also detected, with an alignment length of 1930 aa and being 30% identical to the RdRp of *Xinmovirid*ae family representatives. Extended contigs encoding proteins homologous to RdRp, ORF3, and hypothetical proteins of the *Virgaviridae* family were also identified. Moreover, several contigs have exhibited homology of their protein products to uncharacterized proteins of various representatives of the *Poxviridae, Hytrosaviridae, Partitiviradae, Bornaviridae, Rhabdoviridae, Parvoviridae* families, and several viruses of unknown viral families.

The protein products encoded by the same contig tend to exhibit significant homology with related viruses. To understand how viral annotations are related to each other, we additionally clustered contigs based on the homology of the encoded proteins with proteins from different viral families (Figure 4). The maximum alignment length of the proteins encoded by each contig was calculated for each contig relative to the proteins of each of the presented viral families. The alignment lengths of the proteins encoded by each of the contigs with the amino acid sequences of each of the viral families are presented as values in the heatmap cells. Figure 4 shows the contigs with only a unique homology pattern.

**Figure 4.**
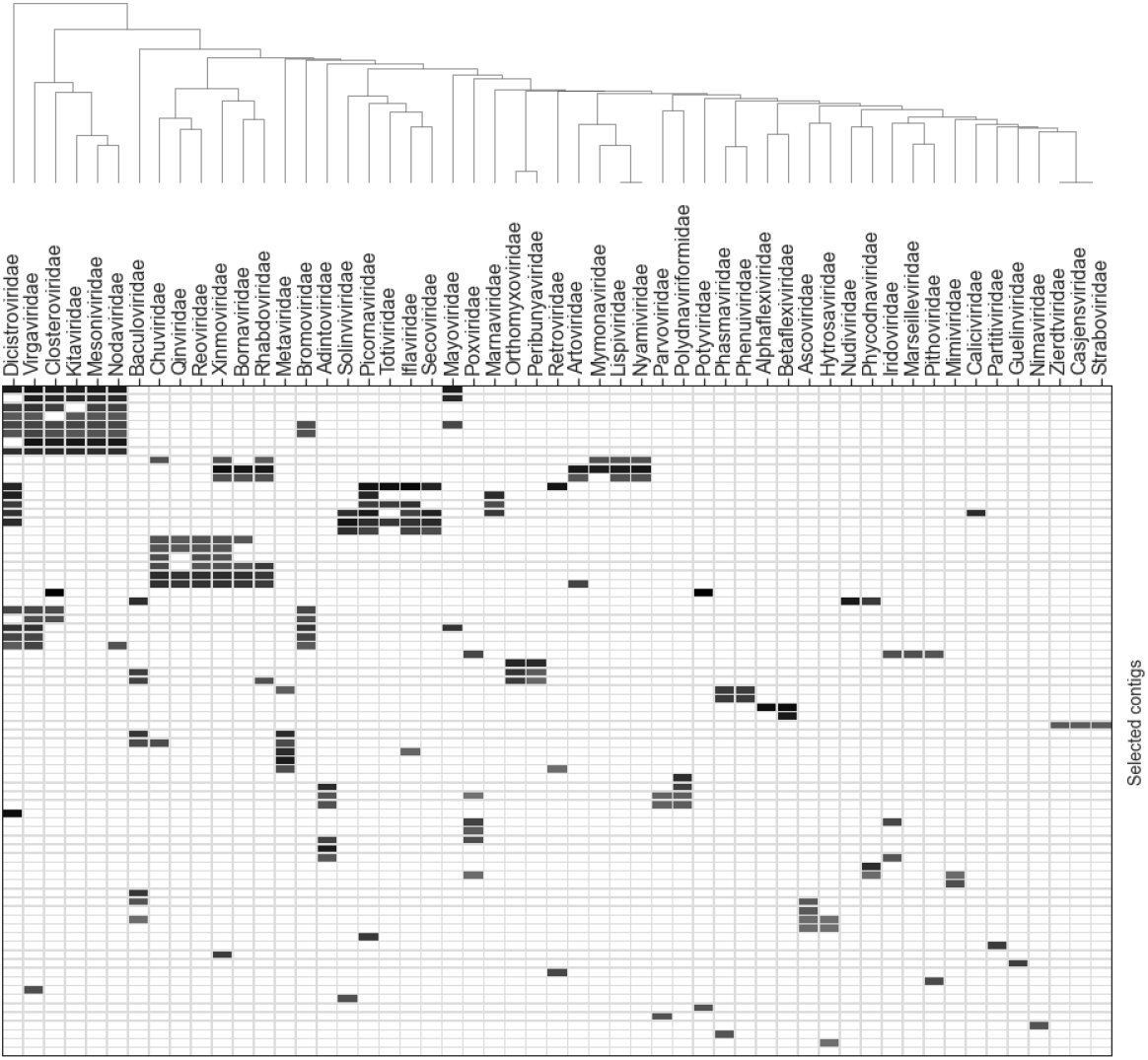
Clustering of selected contigs and viral families based on the homology of contig-encoded proteins with proteins from different virus families.

It can be seen that contigs are clustered largely based on the phylogenetic kinship of viral groups. In particular, contigs with proteins homologous to those of viruses of the *Dicistroviridae* family are clustered with RNA-containing viruses, including other families of the *Picornavirales* order (*Picornaviridae, Iflavirudae, Secoviridae, Caliciviridae, Marnaviridae*), as well as with virus families of the Martellivirales order (*Virgaviridae, Kitaviridae, Closteroviridae*). Contigs encoding proteins that are homologous to proteins of phylogenetically close orders *Mononegavirales (Xinmoviridae, Bornaviridae, Rhabdoviridae*) and *Jingchuvirales (Qinviridae, Chuviridae*) are also clustered. A separate cluster is formed by contigs encoding proteins homologous to those of viruses of the families *Xinmoviridae, Bornaviridae, Rhabdoviridae, Artoviridae, Mymonaviridae, Lispiviridae*, and *Nyamiviridae* of the order *Mononegavirales*. A number of contig clusters encode the proteins that are homologous to proteins of *Nucleocytoviricota* representatives (families *Iridoviridae, Ascoviridae, Phycodnaviridae, Pithoviridae, Mimiviridae, Marseilleviridae*).

A full version of the table containing the data used in Figure 4 is presented in Supplementary Table 8. An interactive visualization of all the contigs analyzed can be found at the project repository at https://github.com/starchevskayamaria17/uncoVir.

Most contigs whose protein products were not classified as viral proteins were found to have homology with the amino acid sequences of arthropods, and mainly with insect proteins.

### Non-viral protein hits are significantly enriched with unknown and uncharacterized proteins as compared to viral protein hits

The next step was to analyze the enrichment of the contigs under study with unknown and uncharacterized proteins. The statistical analysis was performed using Fisher’s exact test for unpaired sets (nonoverlapping sets of contigs) and the McNemar test for paired sets (contigs whose proteins have homologs among both viral and non-viral proteins). The entire set of identified homologous proteins was divided into the following groups: “viral” and “non-viral,” and “known” and “unknown.” Proteins with the following keywords in their name were considered unknown: unnamed, uncharacterized, unknown, or hypothetical. The set of contigs was also categorized into groups: “viral” and “non-viral” contigs whose proteins have and lack homology with viral proteins, respectively (Figure 5).

**Figure 5.**
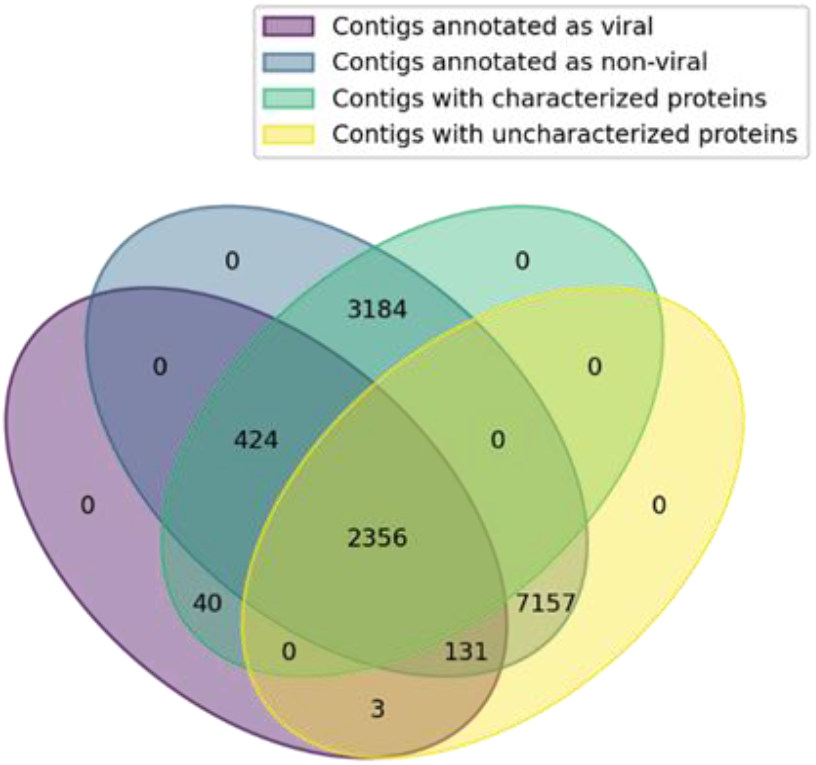
Venn diagram illustrating the distribution of contigs into groups: viral/non-viral and with known/unknown proteins. The number of contigs with “unknown” proteins among non-viral contigs exceeds the number among viral contigs.

The analysis was performed for all “viral” contigs, which were found to be homologous with any of the viral families, and “non-viral” contigs, whose proteins were found to be homologous only with non-viral proteins. A total of 2,817 “known” and 137 “unknown” proteins were found for “viral” contigs, and 3,144 and 7,033 were found for “non-viral” contigs, respectively. Thus, the enrichment with “unknown” proteins among the non-viral proteins was shown to be significant (p < 10^-6^, one-way Fisher’s exact test).

We also performed unknown protein enrichment analysis for contigs with proteins shown to be homologous with viral and non-viral proteins. In this case, 2817 “known” and 137 “unknown” proteins were found among “viral” proteins, and 427 and 2484 were found among “non-viral” proteins, respectively. Thus, the “unknown” protein enrichment among non-viral proteins has been shown to be significant (p < 10^-6^, one-way McNemar test). This finding can be explained by the fact that many eukaryotic genes encoding uncharacterized and hypothetical proteins may be of viral origin.

### Alignment results

The mapping of reads to viral genomes and the extraction of consensus sequences yielded the near full-length or extended genomic sequences for a number of viruses. Shown in Table 2 are the sizes of the reference genomes and the most extended consensus sequences (in nucleotides and % relative to the reference size). The extended information about the alignments is provided in Supplementary Table 9.

**Table 2.** Characteristics of reads alignments to viral genomes. This table specifies the family, genus, and species of the virus, genome coverage information (maximum, average, and median, in nucleotides and percentages), the minimum and maximum genome lengths stored in the BigViralDB database for the specified viral family, and the number of *L. decemlineata* genomic and transcriptomic projects where these viral sequences were found.

| Family | Species | Coverage, bases |  |  | Coverage, % |  |  | Genome length |  | Projects |
| --- | --- | --- | --- | --- | --- | --- | --- | --- | --- | --- |
|  |  | min | max | median | min | max | median | min | max |  |
| Potyviridae | Potato virus Y | 55 | 9561 | 100,50 | 0,57 | 98,63 | 1,03 | 9595 | 9800 | 8 |
| Betaflexiviridae | Potato virus S | 55 | 8481 | 6336,00 | 0,65 | 99,95 | 74,56 | 8453 | 8515 | 10 |
| Dicistroviridae | Aphis glycines virus 3 | 133 | 6691 | 3412,00 | 1,31 | 70,84 | 36,08 | 9445 | 10131 | 2 |
|  | Aphid lethal paralysis virus | 6635 | 6635 | 6635,00 | 67,57 | 67,57 | 67,57 | 9819 | 9819 | 1 |
| Betaflexiviridae | Potato virus H | 6520 | 6520 | 6520,00 | 77,53 | 77,53 | 77,53 | 8410 | 8410 | 1 |
| Unclassified | Wuhan insect virus 33 | 5877 | 5877 | 5877,00 | 57,44 | 57,44 | 57,44 | 10232 | 10232 | 1 |
| Dicistroviridae | Homalodisca coagulata virus 1 | 3998 | 3998 | 3998,00 | 42,78 | 42,78 | 42,78 | 9345 | 9345 | 1 |
| Retroviridae | Human endogenous retrovirus K | 90 | 2908 | 277,00 | 0,95 | 30,70 | 2,92 | 9472 | 9472 | 6 |
| Virgaviridae | Turnip vein-clearing virus | 2795 | 2795 | 2795,00 | 44,29 | 44,29 | 44,29 | 6311 | 6311 | 1 |
| Herpesviridae | Human betaherpesvirus 5 | 50 | 2647 | 191,00 | 0,02 | 1,15 | 0,08 | 229209 | 235732 | 14 |
| Togaviridae | Semliki Forest virus | 100 | 1838 | 622,00 | 0,69 | 12,65 | 4,28 | 14529 | 14529 | 12 |
| Unclassified | Aphis glycines virus 2 | 1567 | 1567 | 1567,00 | 32,31 | 32,31 | 32,31 | 4850 | 4850 | 1 |
|  | Hubei picorna-like virus 15 | 1395 | 1395 | 1395,00 | 14,08 | 14,08 | 14,08 | 9910 | 9910 | 1 |
| Dicistroviridae | Rhopalosiphum padi virus | 1235 | 1235 | 1235,00 | 12,29 | 12,29 | 12,29 | 10045 | 10045 | 1 |
| Unclassified | Wuhan aphid virus 1 | 1185 | 1185 | 1185,00 | 41,89 | 41,89 | 41,89 | 2829 | 2829 | 1 |
| Baculoviridae | Autographa californica multiple nucleopolyhedrovirus | 50 | 1086 | 294,00 | 0,04 | 0,92 | 0,23 | 118582 | 138991 | 8 |
| Dicistroviridae | Big Sioux River virus | 1064 | 1064 | 1064,00 | 10,39 | 10,39 | 10,39 | 10237 | 10237 | 1 |
| Paramyxoviridae | Canine morbillivirus | 105 | 1017 | 453,00 | 0,56 | 5,40 | 2,41 | 18826 | 18826 | 12 |
| Adenoviridae | Human mastadenovirus C | 55 | 775 | 182,00 | 0,15 | 2,16 | 0,51 | 35926 | 35958 | 5 |
| Polydnaviriformidae | Chelonis inanitus bracovirus | 368 | 722 | 545,00 | 1,72 | 3,38 | 2,55 | 21376 | 21376 | 2 |
| Flaviviridae | Pestivirus A | 36 | 401 | 92,50 | 0,25 | 3,16 | 0,68 | 12295 | 15752 | 14 |
| Pospiviroidae | Citrus exocortis viroid | 121 | 376 | 214,00 | 30,56 | 94,95 | 54,04 | 396 | 396 | 13 |
| Polydnaviriformidae | Diolcogaster facetosa bracovirus | 73 | 345 | 101,00 | 0,07 | 0,98 | 0,10 | 12859 | 104039 | 7 |

The sequences representing 99.63% and 99.95% of corresponding reference genomes were obtained for Potato virus Y (family *Potyviridae*) and Potato virus S (family *Betaflexiviridae*), respectively. The size of consensus sequences obtained as a result of the alignment for five members of the family *Dicistrovir*idae ranged from 10.39% to 70.84% of their genome size. Additionally, the alignment allowed us to obtain the consensus sequences of genome fragments of other viruses known to affect insects, such as the *Wuhan insect virus 33*, and the representatives of the families *Virgaviridae, Baculoviridae*, and *Polydnaviriformidae*.

### Analysis of selected coleopteran species reference genomes to detect bracoviral genetic fragments

The k-mer analysis and the contigs annotation revealed numerous sequences belonging to members of the genus *Bracoviriform* (family *Polydnaviriformidae*). The analysis of the reference genome *Leptinotarsa decemlineata* revealed numerous sequences homologous to the genus *Bracoviriform*, with a total length of approximately 25 kbp. Several other coleopteran genomes were also analyzed. Figure 6 shows a heatmap of the distribution of k-mers classified as *Bracoviriform*, depending on the e-value.

**Figure 6.**
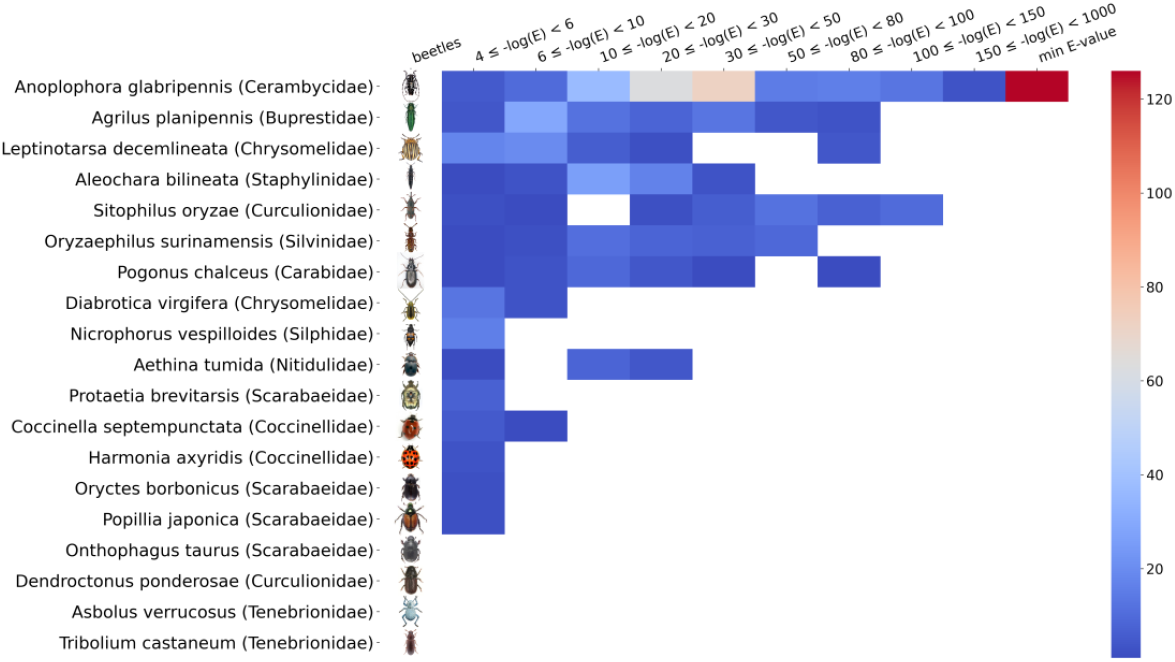
Distribution of bracovirus k-mers depending on e-value in reference genomes of several members of *Coleoptera*, including *Leptinotarsa decemlineata*. The color indicates the number of bracovirus k-mers identified within the specified e-value interval (blastn short). The absence of the color designates the absence of a match.

Two sequences with homology to the segments of the bracovirus genomes were selected for the subsequent experimental studies. The nucleotide sequences of contigs 1513 and 2100 nucleotides long were homologous to *Cotesia sesamiae Mombasa bracovirus* (EF710639.1, the coordinates of the reference sequence alignment: 57046-58557, identity percentage 100%) and *Cotesia vestalis bracovirus* segment 35 (HQ009558.1, the coordinates of the reference sequence alignment: 1797-3455, identity percentage of 98.13%) respectively. These contigs were found to be also aligned to the *Leptinotarsa decemlineata* genome. The first bracovirus contig was aligned to NW_019290920.1, and the coordinates of the reference sequence alignment were 2663-4169; the second contig was aligned to NW_019296350.1, and the coordinates of the reference sequence alignment were 9973-11595. The alignment allows us to assume the insertion of viral DNA of *Bracoviriform* genus into the genome of *Leptinotarsa decemlineata*.

We calculated the primers so as to confirm the presence of bracovirus fragments in the genetic material of Leptinotarsa decemlineata, to prove the insertion into the Leptinotarsa decemlineata genome, and to exclude the presence of fragments containing regions flanking the insertion in the corresponding bracovirus genome. The PCR analysis of DNA obtained from different tissues of *Leptinotarsa decemlineata* confirmed the presence of both bracovirus fragments and revealed the absence of fragments containing the regions flanking the insertion in the bracovirus genome. The nucleotide sequence of the fragments was confirmed by Sanger sequencing. The insertion into the *Leptinotarsa decemlineata* genome was not confirmed by this experiment, which may be due to assembly errors or high variability of corresponding regions of the *L. decemlineata* genome (Supplementary figures SF3-6).

## 4. Discussion

Among the annotated sequences, contigs of viral origin can be divided into several groups: homologous to genetic sequences of insect viruses (representatives of the families *Bornaviridae, Caliciviridae, Dicistroviridae, Iflaviridae, Iridoviridae, Orthomyxoviridae, Partitiviridae, Picornaviridae, Poxviridae, Virgaviridae, Xinmoviridae*), plant viruses (representatives of families *Betaflexiviridae, Bromoviridae, Kitaviridae, Potyviridae*), endogenous retroviral elements (*Retroviridae, Metaviridae*), endogenous viral elements of non-retroviral origin (*Adintoviridae, Ascoviridae, Chuviridae, Hytrosaviridae, Nimaviridae, Nudiviridae, Phasmaviridae, Polydnaviriformidae, Rhabdoviridae, Baculoviridae*), bacteriophage sequences and protozoan viruses (not described within this study) and numerous viral sequences of unknown family primarily affecting insects.

### Insect viruses

The numerous sequences of insect-infecting viruses were found in the data analyzed. The special attention should be given to the viruses of the families *Iflaviridae* and *Dicistroviridae* (the members of the order *Picornavirales*), whose genetic sequences were detected at different stages of the analysis. The members of the families in question are characterized by a rother small ss(+)RNA genome measuring 8–10 kb for the family *Dicistroviridae* and 9–11 kb for the family *Iflaviridae*. It is worth noting that new virus species of these families are often identified by metagenomic analysis in NGS data of various insects, particularly mosquitoes [27]. The *Homalodisca coagulata dicistrovirus* also identified in this way is the virus of a glassy-winged sharpshooter *Homalodisca coagulata* (Cicadellidae), which feeds on hundreds of plant species and carries plant pathogens, causing enormous damage to agriculture [28]. Here we have obtained numerous sequences attributed to these two families of viruses, with contig lengths commensurating with their complete genome lengths, some of which are probably the genetic sequences of new viruses of these families. The contigs annotated as homologous to *Lampyris noctiluca iflavirus 1* reach a length of 10 kb. However, the amino acid sequence of the polyprotein encoded by these contigs is only 30% identical to that of the iflavirus. In this study, we have identified the sequences homologous to *Homalodisca coagulata dicistrovirus* (comprising over 40% of the genome), *Aphid lethal paralysis* virus (up to 70% of the genome and an extended contig with a length of 3468 nucleotides, encoding a polypeptide that is 92% identical to the viral protein), *Aphid glycines virus 3* (over 70% of the genome), *Aphis gossypii* virus (a 6,635-nucleotide-long extended contig encoding a protein product that is 93% identical to the viral polyprotein), and others. The majority of known members of the family *Dicistroviridae* have been shown to infect only a limited number of related insect species (belonging to the same order). However, there are viruses in this family that infect a vast range of hosts. For example, *Cricket paralysis virus* can infect more than 20 insect species from different orders [29].

Earlier studies suggested that the Colorado potato beetle could be infected by viruses of the *Baculoviridae* and *Iridoviridae* families [30]. Various baculoviruses are known to affect a wide variety of insect taxonomic groups. Currently, the members of the family *Baculoviridae* are used as safe and effective biopesticides to protect forests, fields, and horticultural crops against lepidopteran pests [31]. Most of the contigs found in the genomic data and annotated as baculoviruses exhibit homology to the F-protein gene of various members of the genera *Alphabaculovirus* and *Betabaculovirus* and will be described in more detail below in the “Endogenous viral elements” section. Although, a number of contigs are found to be homologous with baculovirus sequences different from the F-protein gene, the low length of such fragments does not allow us to annotate them reliably.

Among the members of the Nucleocytoviricota type, a number of viruses of the families Iridoviridae, Poxviridae, and Ascoviridae are known to affect insects. However, despite the presence of contigs assigned to these families, the low percentage of identity (30–40%) and alignment length (64 to 191 aa), as well as the short length of contigs (from 500 to 2044 nucleotides) do not allow conclusions to be drawn about the presence of genetic material of specific viruses of these families in the samples analyzed. These sequences may also be fragments of endogenous viral elements integrated into the *L. decemlineata* genome over the course of evolution. The presence of endogenous viral elements of iridovirus origin in insect genomes is indicated, in particular, by the fact that some representatives of leaf-feeding beetles, such as *Gastrophysa atrocyanea*, were shown to produce a diapause-specific peptide that acts as a blocker of potential-dependent calcium channels and provides antifungal activity and is homologous to the sequence of the peptide encoded by *Iridoviridae*. The sequences encoding these peptides have been suggested to be suitable as probes for analyzing insects and this group of viruses [32].

For several other viruses affecting insects, such as members of the families *Ortomyxoviridae, Virgaviridae, Xinmoviridae, Partiviridae*, and viruses with unspecified families, such as *Wuhan insect virus 33, Hubei picorna-like virus 15, Allermuir Hill virus 1*, the extended amino acid alignments with various structural and nonstructural viral proteins have been found, with corresponding virus-positive contigs being highly extended.

### Plant viruses

Plant pathogenic viruses pose a serious threat to livestock, agricultural, horticultural and ornamental crops. *Potato virus Y* detected in this study belongs to the RNA-containing virus family *Potyvirus* of the family *Potyviridae*, which represents approximately 30% of all known plant pathogenic viruses. *Potato virus Y*, usually transmitted by aphids, can significantly reduce the potato yields [4]. *Potato virus S*, the full-length genome of which was also identified in this work, belongs to the RNA-containing viruses of the *Betaflexviridae family*. Despite having less pronounced disease symptoms in the affected plants, *Potato virus S* can destroy up to 20% of the crops [33]. These results confirm the presence of vast amounts of potato viruses in *L. decemlineata* tissues and especially in gut and head samples.

In addition, we identified 63 extended (up to 8.5 kb) contigs belonging to the *Papaya mottle associated virus*, with their protein products being homologous to RdRp. Additionally, we identified several plant viruses positive contigs with protein products being homologous to those of *Garlic common virus*, *Bromoviridae*, and *Kitaviridae* with alignment length less than 200 aa and identity less than 50%. Thus, it’s hardly possible to make any strong conclusions on these contigs.

### Endogenous virus elements

The ability of many viruses to integrate their genetic material into the host genome is widely known. It was reported that 0.59% of the genome of *L. decemlineata* contains LTR regions, including Copia, Gypsy, Gypsy-Cigr, and Pao [7]. This study has identified numerous retroviral sequences homologous to the members of the families *Retroviridae* and, to a greater extent, *Metaviridae*, characteristic to insects (LTR retrotransposon, Ty3/gypsy family), as well as sequences homologous to unclassified viruses AAToV1 and AAToV2. Along with integrated retroviral sequences, we have discovered viral sequences of non-retroviral origin, called endogenous viral elements (EVEs). The currently known insect EVEs belong to at least 28 viral families [34]. Endogenization of viral sequences into the host genome and EVE formation can occur via nonhomologous recombination or with participation of reverse transcriptase and integrase of endogenous retrotransposons. EVEs can also contribute to the emergence of new functional host genes and even play an antiviral role by generating small RNA and thereby preventing the maturation of viral particles at various stages of the viral life cycle [35–37].

Recent studies of insect genomes have uncovered numerous genetic sequences of negative sense RNA-containing viruses [34,38]. Such studies confirm the fact that arthropods are one of the main reservoirs of viral genetic diversity and undoubtedly play an essential role in virus evolution. Genetic sequences of the members of the family *Chuviridae*, characterized by an unusual ordering of genes, have been found in many genomic sequences of various invertebrates [39]. In particular, integrated genes of *Chuviridae* glycoproteins were found in the genomes of *Dendroctonus ponderosae* and *Tribolium castaneum* (*Coleoptera*) [38]. For the mosquitoes *Aedes aegypti* and *Aedes albopictus*, the presence of endogenous Chuviridae-like genes in Bel/Pao elements (family *Belpaoviridae*) was described. In our study in genomic data samples, we have found contigs encoding proteins similar to glycoproteins of viruses of the *Chuviridae* family, including the *Coleopteran chu-related virus*. In addition, in the genomic data of *L. decemlineata*, we have also found sequences homologous to the genes of viruses of the families *Phasmaviridae, Rhabdoviridae, Xinmoviridae, Bornaviridae*, and *Orthomyxoviridae*. These viruses have previously been shown to be present as EVEs within the genomes of various insect families [34].

Of interest is the fact that sequences homologous to the baculovirus F-protein gene were found in the genomic data of *L. decemlineata*. It is believed that the ancestral baculoviruses could replicate only in the midgut epithelial cells of insects, and the F-protein gene acquired during the evolution allowed the virus to penetrate the hemocoel and cause generalized infection. Previously, the orthologs of the baculovirus F-protein gene were found in the endogenous retroviral sequences of lepidopterans, dipterans, and other insect species, indicating that the regions encoding F-protein homologs identified in our study can also be incorporated into the genome of *L. decemlineata* [40,41] and thus may indicate that the Colorado potato beetle can also be affected by baculoviruses.

We also have identified numerous extended fragments homologous to the genetic sequences of viruses of the genus *Bracoviriform*. The members of the genus *Bracoviriform* belong to the family *Polydnaviriformidae* and are associated with parasitoid wasps, including approximately 18,000 species of five subfamilies of the family Braconidae (*Microgastrinae, Cardiochilinae, Miracinae, Khoikholinae, and Cheloninae*). The genomes of *Polydnaviriformidae* family viruses are segmented. Their segments have multiple copies and present in virions non-equimolarly in the form of circular supercoiled double-stranded DNA measuring from 2 to 31 kbp. The cumulative non-redundant size of the bracovirus genome varies from 190 to 500 kbp [42]. The viral genome is transmitted as proviral DNA in parasitoid wasp cells and circular episomal DNA within virions. Replication of the virus genome and virion assembly occur in ovarian cells, with viral particles being released and accumulated in the lumen of the oviduct and entering the host larvae or eggs during wasps laying their own eggs. In other wasp cells, the virus does not replicate and exists predominantly in a proviral form. Interestingly, the expression of viral genes causes changes in the physiology of the infected host and suppresses the encapsulation of wasp eggs, promoting the development and emergence of wasps [43]. Wasps parasitize on the larvae and eggs of lepidopterans, coleopterans, and hymenopterans. Despite the absence of replication in the cells of lepidopterans, the integration of Bracovirus genome segments has been demonstrated both *in vitro*, in cell cultures, and *in vivo* [42]. Some bracovirus genes were shown to contribute to protecting insects against baculoviruses – lethal pathogens, which explains why these genes have been conserved in the genomes of many lepidopterans [44].

We have found the extended bracovirus sequences in the reference genome of *L. decemlineata* and in the genomes of several other representatives of *Coleoptera*. We have also demonstrated the presence of bracovirus sequences in the genetic material isolated not only from imago and larva insect tissues but also from sterile eggs of *L. decemlineata*, suggesting the integration of bracovirus genomic sequences within the Colorado potato beetle genome. To our knowledge, we are the first to report on the detection of bracovirus sequences in the genomes of the Colorado potato beetle and leaf-feeding beetles. The presence of these fragments in the Colorado potato beetle genome also seems to be associated with parasitoid wasps, such as the wasp *Edovium puttleri*, which parasitizes the eggs of *Leptinotarsa decemlineata* [45].

Besides, the other sequences identified in our study, which are homologous to the genetic sequences of *Nudiviridae* family viruses, are also probably associated with bracovirus genetic fragments, given that the genus *Bracoviriform* originated from the *Nudiviridae* family [46].

Thus, the metagenomic studies of accumulated public high-throughput sequencing data serve as an important source of information for the search for new viruses. This work highlights the significance of metagenomic studies of the genomic and transcriptomic data obtained from insect pests for a better understanding of their ecology and evolution. The discovery of genetic sequences potentially belonging to insect-infecting viruses points to the possibility of expanding the arsenal of biological methods to control the Colorado potato beetle population based on entomopathogenic viruses.

## Author Contributions

Conceptualization, D.A. and Y.V.; methodology, M.S., D.A. and Y.V.; scripts and analytical pipelines, visualization, resources and data curation, M.S. and D.A.; biological samples collection, U.R., O.P. and V.K.; DNA extraction, PCR and sequencing, T.T., T.B. and S.B.; writing—original draft preparation, M.S. and D.A.; writing—review and editing, M.S., D.A., U.R. and V.K.; supervision, D.A.; funding acquisition, D.A. and V.K. All authors have read and agreed to the published version of the manuscript.

## Funding

This work was supported by the Ministry of Science and Higher Education of Russian Federation (agreement # 075-15-2019-1665). Insect collection and tissue sample preparation were supported by the Russian Scientific Foundation (grant # 22-14-00309).

## Data Availability Statement

All links to publicly archived datasets analyzed during the study are provided in “Materials and Methods” section. Additional data is provided in Supplementary Materials. The analytical pipeline, examples and jupyther notebooks are freely available at https://github.com/starchevskayamaria17/uncoVir

## Conflicts of Interest

The authors declare no conflict of interest. The funders had no role in the design of the study; in the collection, analyses, or interpretation of data; in the writing of the manuscript; or in the decision to publish the results.

## Notes

https://github.com/starchevskayamaria17/uncoVir

### Competing Interest Statement

The authors have declared no competing interest.

## References

1. Rosario, K.; Breitbart, M. Exploring the Viral World through Metagenomics. Curr. Opin. Virol. 2011, 1, 289–297, doi:10.1016/j.coviro.2011.06.0041.

2. Varghese, F.S.; van Rij, R.P. Insect Virus Discovery by Metagenomic and Cell Culture-Based Approaches. Methods Mol. Biol. 2018, 1746, 197–213, doi:10.1007/978-1-4939-7683-6_16.

3. Cingel, A.; Savić, J.; Lazarević, J.; Ćosić, T.; Raspor, M.; Smigocki, A.; Ninković, S. Extraordinary Adaptive Plasticity of Colorado Potato Beetle: “Ten-Striped Spearman” in the Era of Biotechnologicalwarfare. Int. J. Mol. Sci. 2016, 17, doi:10.3390/ijms17091538.

4. Petek, M.; Rotter, A.; Kogovšek, P.; Baebler, S.; Mithöfer, A.; Gruden, K. Potato Virus Y Infection Hinders Potato Defence Response and Renders Plants More Vulnerable to Colorado Potato Beetle Attack. Mol. Ecol. 2014, 23, 5378–5391, doi:10.1111/mec.12932.

5. Sablon, L.; Dickens, J.C.; Haubruge, É.; Verheggen, F.J. Chemical Ecology of the Colorado Potato Beetle, Leptinotarsa Decemlineata (Say) (Coleoptera: Chrysomelidae), and Potential for Alternative Control Methods. Insects 2012, 4, 31–54, doi:10.3390/insects4010031.

6. Alyokhin, A.; Baker, M.; Mota-Sanchez, D.; Dively, G.; Grafius, E. Colorado Potato Beetle Resistance to Insecticides. Am. J. Potato Res. 2008, 85, 395–413, doi:10.1007/s12230-008-9052-0.

7. Schoville, S.D.; Chen, Y.H.; Andersson, M.N.; Benoit, J.B.; Bhandari, A.; Bowsher, J.H.; Brevik, K.; Cappelle, K.; Chen, M.- J.M.; Childers, A.K.; et al. A Model Species for Agricultural Pest Genomics: The Genome of the Colorado Potato Beetle, Leptinotarsa Decemlineata (Coleoptera: Chrysomelidae). Sci. Rep. 2018, 8, 1931, doi:10.1038/s41598-018-20154-1.

8. Tanwar, R.S.; Dureja, P.; Rathore, H.S. Biopesticides; Elsevier Inc., 2012; ISBN 9781439836255.

9. Alyokhin, A.; Kryukov, V. Chapter 25 - Ecology of a Potato Field. In; Alyokhin, A., Rondon, S.I., Gao, Y.B.T.-I.P. of P. (Second E., Eds.; Academic Press, 2022; pp. 451–462 ISBN 978-0-12-821237-0.

10. Yu, Y.; Yuan, Y.; Gao, M. Construction of an Environmental Safe Bacillus Thuringiensis Engineered Strain against Coleoptera. Appl. Microbiol. Biotechnol. 2016, 100, 4027–4034, doi:10.1007/s00253-015-7250-5.

11. Beas-Catena, A.; Sánchez-Mirón, A.; García-Camacho, F.; Contreras-Gómez, A.; Molina-Grima, E. Baculovirus Biopesticides: An Overview. J. Anim. Plant Sci. 2014, 24, 362–373.

12. Jones, R.A.C. Plant and Insect Viruses in Managed and Natural Environments: Novel and Neglected Transmission Pathways; 1st ed.; Elsevier Inc., 2018; Vol. 101;.

13. Selman, B.J. Viruses and Chrysomelidae. Biol. Chrysomelidae 1988, 379–387, doi:10.1007/978-94-009-3105-3_21.

14. Kumar, A.; Congiu, L.; Lindström, L.; Piiroinen, S.; Vidotto, M.; Grapputo, A. Sequencing, de Novo Assembly and Annotation of the Colorado Potato Beetle, Leptinotarsa Decemlineata, Transcriptome. PLoS One 2014, 9, doi:10.1371/journal.pone.0086012.

15. M. Kudla, K. Gutowska, J. Synak, M. Weber, K.S. Bohnsack, P. Lukasiak, T. Villmann, J. Blazewicz, M.S. Virxicon: A Lexicon of Viral Sequences. Bioinformatics 2020, 36, 5507–5513.

16. Gleizes, A.; Laubscher, F.; Guex, N.; Iseli, C.; Junier, T.; Cordey, S.; Fellay, J.; Xenarios, I.; Kaiser, L.; Le Mercier, P. Virosaurus a Reference to Explore and Capture Virus Genetic Diversity. Viruses 2020, 12, 1–13, doi:10.3390/v12111248.

17. Mölder, F.; Jablonski, K.P.; Letcher, B.; Hall, M.B.; Tomkins-tinch, C.H.; Sochat, V.; Forster, J.; Lee, S.; Twardziok, S.O.; Kanitz, A.; et al. Sustainable Data Analysis with Snakemake. F1000Research 2021, 1–25.

18. Bolger, A.M.; Lohse, M.; Usadel, B. Trimmomatic: A Flexible Trimmer for Illumina Sequence Data. Bioinformatics 2014, 30, 2114–2120, doi:10.1093/bioinformatics/btu170.

19. Langmead, B.; Salzberg, S.L. Fast Gapped-Read Alignment with Bowtie 2. Nat. Methods 2012, 9, 357–359, doi:10.1038/nmeth.1923.

20. Wood, D.E.; Salzberg, S.L. Kraken: Ultrafast Metagenomic Sequence Classification Using Exact Alignments. Genome Biol. 2014, 15, R46, doi:10.1186/gb-2014-15-3-r46.

21. Breitwieser, F.P.; Salzberg, S.L. Pavian: Interactive Analysis of Metagenomics Data for Microbiome Studies and Pathogen Identification. Bioinformatics 2020, 36, 1303–1304, doi:10.1093/bioinformatics/btz715.

22. Bankevich, A.; Nurk, S.; Antipov, D.; Gurevich, A.A.; Dvorkin, M.; Kulikov, A.S.; Lesin, V.M.; Nikolenko, S.I.; Pham, S.; Prjibelski, A.D.; et al. SPAdes: A New Genome Assembly Algorithm and Its Applications to Single-Cell Sequencing. J. Comput. Biol. 2012, 19, 455–477, doi:10.1089/cmb.2012.0021.

23. Rotskaya, U.N.; Kryukov, V.Y.; Kosman, E.; Tyurin, M.; Glupov, V. V. Identification of the Ricin-b-Lectin Ldrblk in the Colorado Potato Beetle and an Analysis of Its Expression in Response to Fungal Infections. J. Fungi 2021, 7,doi:10.3390/jof7050364.

24. Kryukov, V.Y.; Kabilov, M.R.; Smirnova, N.; Tomilova, O.G.; Tyurin, M. V.; Akhanaev, Y.B.; Polenogova, O. V.; Danilov, V.P.; Zhangissina, S.K.; Alikina, T.; et al. Bacterial Decomposition of Insects Post-Metarhizium Infection: Possible Influence on Plant Growth. Fungal Biol. 2019, 123, 927–935, doi:10.1016/j.funbio.2019.09.012.

25. Polenogova, O. V.; Noskov, Y.A.; Yaroslavtseva, O.N.; Kryukova, N.A.; Alikina, T.; Klementeva, T.N.; Andrejeva, J.; Khodyrev, V.P.; Kabilov, M.R.; Kryukov, V.Y.; et al. Influence of Bacillus Thuringiensis and Avermectins on Gut Physiology and Microbiota in Colorado Potato Beetle: Impact of Enterobacteria on Susceptibility to Insecticides. PLoS One 2021, 16, 1–22, doi:10.1371/journal.pone.0248704.

26. Untergasser, A.; Cutcutache, I.; Koressaar, T.; Ye, J.; Faircloth, B.C.; Remm, M.; Rozen, S.G. Primer3-New Capabilities and Interfaces. Nucleic Acids Res. 2012, 40, 1–12, doi:10.1093/nar/gks596.

27. Cholleti, H.; Hayer, J.; Fafetine, J.; Berg, M.; Blomström, A.L. Genetic Characterization of a Novel Picorna-like Virus in Culex Spp. Mosquitoes from Mozambique. Virol. J. 2018, 15, 1–10, doi:10.1186/s12985-018-0981-z.

28. Hunnicutt, L.E.; Mozoruk, J.; Hunter, W.B.; Crosslin, J.M.; Cave, R.D.; Powell, C.A. Prevalence and Natural Host Range of Homalodisca Coagulata Virus-1 (HoCV-1). Arch. Virol. 2008, 153, 61–67, doi:10.1007/s00705-007-1066-2.

29. Reinganum, C. The Isolation of Cricket Paralysis Virus from the Emperor Gum Moth, Antheraea Eucalypti Scott, and Its Infectivity towards a Range of Insect Species. Intervirology 1975, 5, 97–102, doi:10.1159/000149886.

30. Hough-Goldstein, J.A.; Heimpel, G.E.; Bechmann, H.E.; Mason, C.E. Arthropod Natural Enemies of the Colorado Potato Beetle. Crop Prot. 1993, 12, 324–334, doi:10.1016/0261-2194(93)90074-S.

31. Bonning, B.C.; Nusawardani, T. Introduction to the Use of Baculoviruses as Biological Insecticides. Methods Mol. Biol. 2007, 388, 359–366, doi:10.1007/978-1-59745-457-5_18.

32. Tanaka, H.; Sato, K.; Saito, Y.; Yamashita, T.; Agoh, M.; Okunishi, J.; Tachikawa, E.; Suzuki, K. Insect Diapause-Specific Peptide from the Leaf Beetle Has Consensus with a Putative Iridovirus Peptide. Peptides 2003, 24, 1327–1333, doi:10.1016/j.peptides.2003.07.021.

33. Ristić, D.; Vučurović, I.; Kuzmanović, S.; Pfaf-Dolovac, E.; Aleksić, G.; Vučurović, A.; Starović, M. The Incidence and Genetic Diversity of Potato Virus S in Serbian Seed Potato Crops. Potato Res. 2019, 62, 31–46, doi:10.1007/s11540-018-9395-y.

34. Gilbert, C.; Belliardo, C. The Diversity of Endogenous Viral Elements in Insects. Curr. Opin. insect Sci. 2022, 49, 48–55, doi:10.1016/j.cois.2021.11.007.

35. Palatini, U.; Contreras, C.A.; Gasmi, L.; Bonizzoni, M. Endogenous Viral Elements in Mosquito Genomes: Current Knowledge and Outstanding Questions. Curr. Opin. Insect Sci. 2022, 49, 22–30, doi:10.1016/j.cois.2021.10.007.

36. Palatini, U.; Miesen, P.; Carballar-Lejarazu, R.; Ometto, L.; Rizzo, E.; Tu, Z.; van Rij, R.P.; Bonizzoni, M. Comparative Genomics Shows That Viral Integrations Are Abundant and Express PiRNAs in the Arboviral Vectors Aedes Aegypti and Aedes Albopictus. BMC Genomics 2017, 18, 1–15, doi:10.1186/s12864-017-3903-3.

37. Wallau, G.L. RNA Virus EVEs in Insect Genomes. Curr. Opin. Insect Sci. 2022, 49, 42–47, doi:10.1016/j.cois.2021.11.005.

38. Li, C.X.; Shi, M.; Tian, J.H.; Lin, X.D.; Kang, Y.J.; Chen, L.J.; Qin, X.C.; Xu, J.; Holmes, E.C.; Zhang, Y.Z. Unprecedented Genomic Diversity of RNA Viruses in Arthropods Reveals the Ancestry of Negative-Sense RNA Viruses. Elife 2015, 2015, 1–26, doi:10.7554/eLife.05378.

39. Di Paola, N.; Dheilly, N.M.; Junglen, S.; Paraskevopoulou, S.; Postler, T.S.; Shi, M.; Kuhn, J.H. Jingchuvirales: A New Taxonomical Framework for a Rapidly Expanding Order of Unusual Monjiviricete Viruses Broadly Distributed among Arthropod Subphyla. Appl. Environ. Microbiol. 2022, 88, doi:10.1128/aem.01954-21.

40. Lung, O.; Blissard, G.W. A Cellular Drosophila Melanogaster Protein with Similarity to Baculovirus F Envelope Fusion Proteins. J. Virol. 2005, 79, 7979–7989, doi:10.1128/jvi.79.13.7979-7989.2005.

41. Rohrmann, G.F.; Karplus, P.A. Relatedness of Baculovirus and Gypsy Retrotransposon Envelope Proteins. BMC Evol. Biol. 2001, 1, doi:10.1186/1471-2148-1-1.

42. Strand, M.R. and Drezen, J.-M. Family Polydnaviridae. 2012, doi:10.1016/B978-0-12-384684-6.00026-4.

43. Strand, M.R.; Burke, G.R. Polydnaviruses: From Discovery to Current Insights. Virology 2015, 479-480, 393–402, doi:10.1016/j.virol.2015.01.018.

44. Gasmi, L.; Boulain, H.; Gauthier, J.; Hua-van, A.; Musset, K.; Jakubowska, A.K.; Aury, J.; Volkoff, A.; Huguet, E. Recurrent Domestication by Lepidoptera of Genes from Their Parasites Mediated by Bracoviruses. 2015, 1–32, doi:10.1371/journal.pgen.1005470.

45. Jansson, R.K.; Groden, E.; Zoology, E.; Experiment, A.; Brunswick, N. PARASITISM OF- LEPTINOTARSA DECEMLINEATA [COLEOPTERA: CHR YSOMELIDAE] BY EDOVUM PUTTLERI [HYMENOPTERA - EULOPHIDAE] IN DIFFERENT CULTIVARS OF EGGPLANT. 1987, 32, 503–510.

46. Wang, Y.; Jehle, J.A. Nudiviruses and Other Large, Double-Stranded Circular DNA Viruses of Invertebrates: New Insights on an Old Topic. J. Invertebr. Pathol. 2009, 101, 187–193, doi:10.1016/j.jip.2009.03.013.

