## Supplementary tables and figures for "The metagenomic analysis of viral diversity in Colorado potato beetle public NGS data": Starchevskaya_Ldec_SupplementaryFig_bioRxiv.docx

**Supplementary Figures**

**SF1.** Distribution of contigs by non-viral families in genomic (top) and transcriptomic (bottom) samples. Total number of annotated contigs: 13 314.

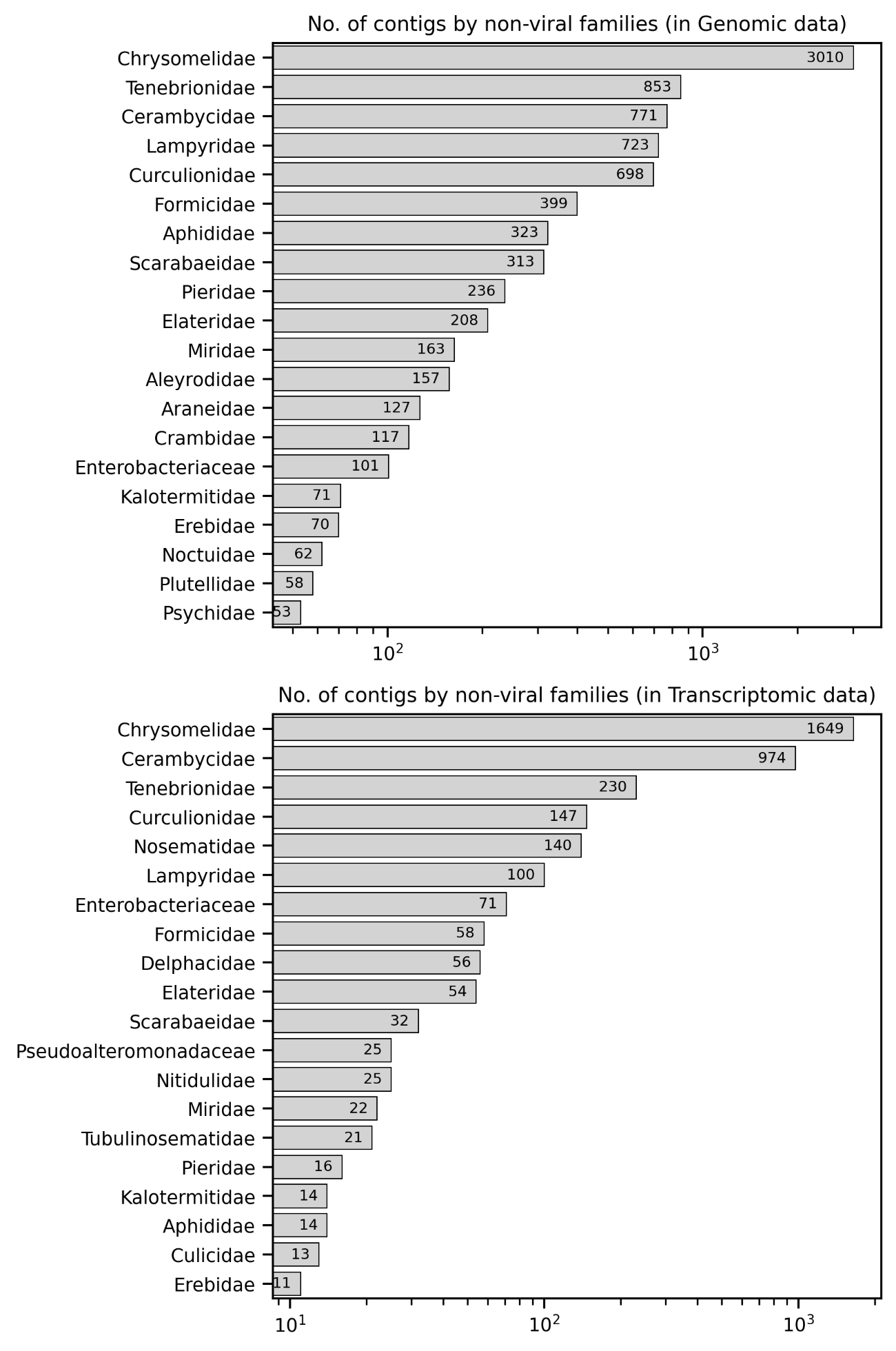

**SF2.** Distribution of contigs by non-viral classes in genomic (top) and transcriptomic (bottom) samples. Total number of annotated contigs: 13 314.

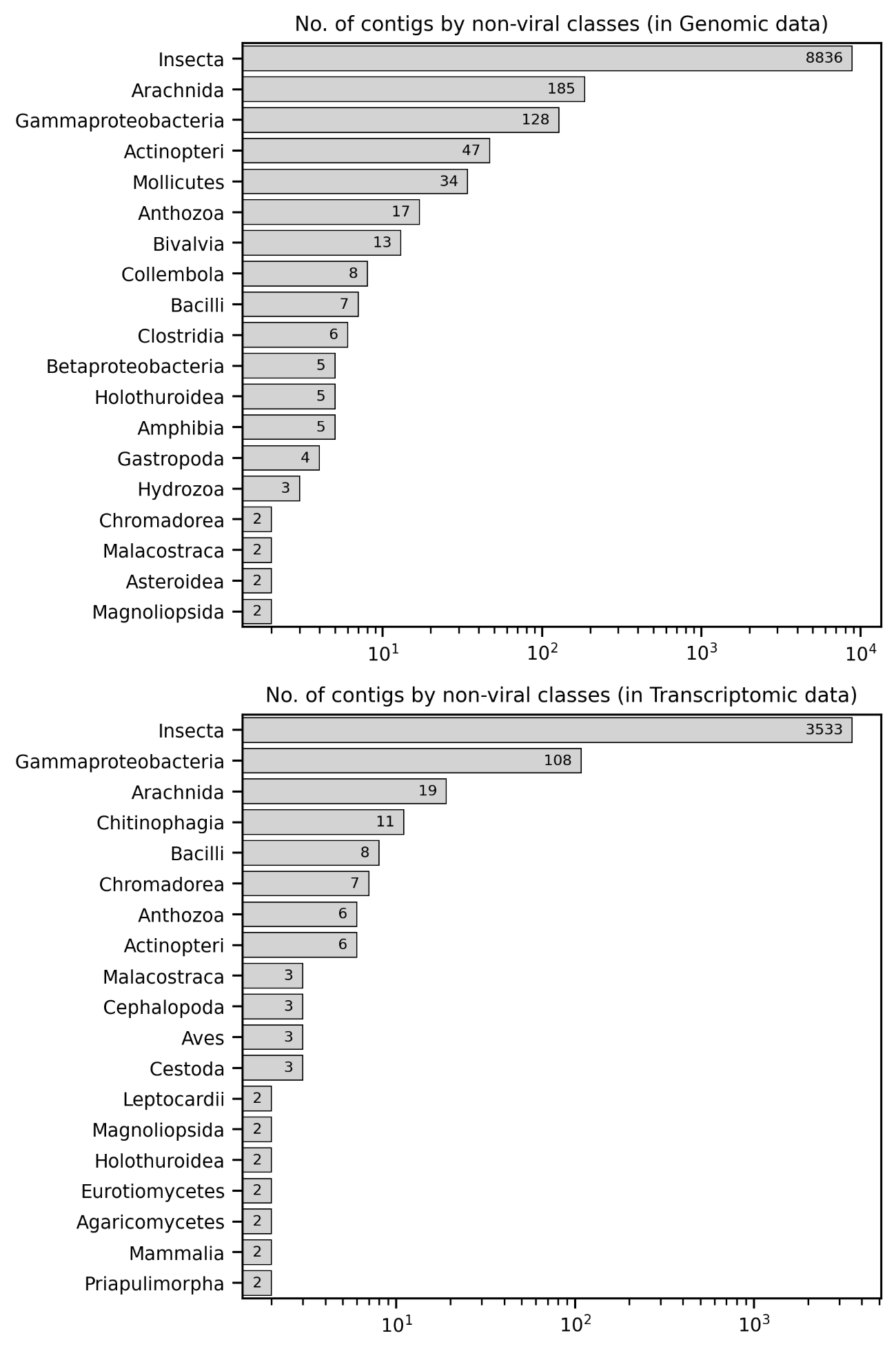

**SF3.** Gel electrophoresis: PCR products showing the presence of the first bracoviral fragment in Colorado potato beetle tissues (primer pairs 1-4)

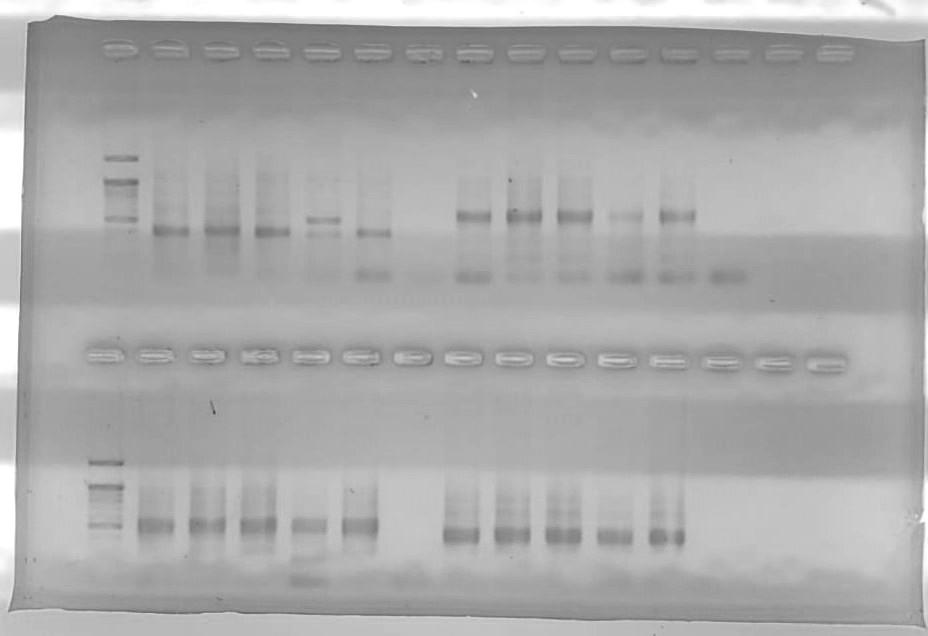

|  | **Br1_F1-Br1_R1 (403 nt)** | | | | | | **Br1_F2_1-Br1_R2_1 (545 nt)** | | | | | |
| --- | --- | --- | --- | --- | --- | --- | --- | --- | --- | --- | --- | --- |
| M | HL | E | PE | I | C | N | HL | E | PE | I | C | N |
|  | **Br1_F2_2-Br1_R2_2 (467 nt)** | | | | | | **Br1_F3-Br1_R3 (380 nt)** | | | | | |
| M | HL | E | PE | I | C | N | HL | E | PE | I | C | N |

M – marker

HL – hemolymph

E – eggs

PE – pure eggs

I – imago

C – cuticle

N – negative control

**SF4.** Gel electrophoresis: PCR products showing the presence of the second bracoviral fragment in Colorado potato beetle tissues (primer pairs 11-15)

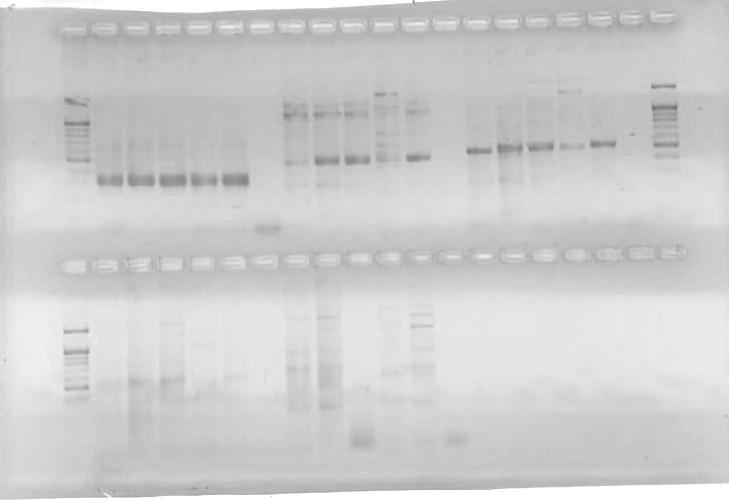

|  | **Br2_F1-Br2_R1**  **(301 nt)** | | | | | | **Br2_F2_1-Br2_R2_1**  **(456 nt)** | | | | | | | **Br2_F2_2-Br2_R2_2**  **(534 nt)** | | | | | |  |
| --- | --- | --- | --- | --- | --- | --- | --- | --- | --- | --- | --- | --- | --- | --- | --- | --- | --- | --- | --- | --- |
| M | HL | E | PE | I | C | N | HL | E | PE | I | | C | N | HL | E | PE | I | C | N | M |
|  | **Br2_F2_3-Br2_R3**  **(519 nt)** | | | | | | **Br2_F3-Br2_R3**  **(243 nt)** | | | | | | |  | | | | | | |
| M | HL | E | PE | I | C | N | HL | E | PE | I | C | | N |  | | | | | | |

M – marker

HL – hemolymph

E – eggs

PE – pure eggs

I – imago

C – cuticle

N – negative control

**SF5.** Gel electrophoresis: PCR products showing no insertion of the first bracoviral fragment into the Colorado potato beetle or bracovirus genome (primer pairs 5-10)

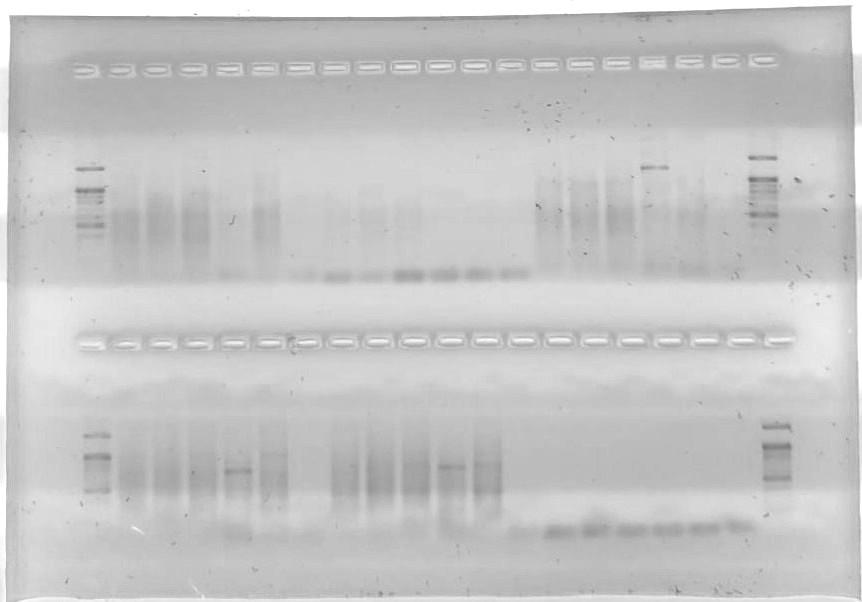

|  | Br1Br_F1-Br1_ins_R1_1  (309 nt) | | | | | | Br1Br_F1-Br1_ins_R1_2  (507 nt) | | | | | | | Br1_ins_F2-Br1Br_R2  (370 nt) | | | | | | | | | |  |
| --- | --- | --- | --- | --- | --- | --- | --- | --- | --- | --- | --- | --- | --- | --- | --- | --- | --- | --- | --- | --- | --- | --- | --- | --- |
| M | HL | E | PE | I | C | N | HL | E | PE | I | | C | N | HL | E | PE | | I | | C | | N | | M |
|  | Br1LD_F1-Br1_ins_R1_1  (401 nt) | | | | | | Br1LD_F1-Br1_ins_R1_2  (599 nt) | | | | | | | Br1_ins_F2-Br1LD_R2  (343 nt) | | | | | | | | | |  |
| M | HL | E | PE | I | C | N | HL | E | PE | I | C | | N | HL | E | | PE | | I | | C | | N | M |

M – marker

HL – hemolymph

E – eggs

PE – pure eggs

I – imago

C – cuticle

N – negative control

**SF6.** Gel electrophoresis: PCR products showing no insertion of the second bracoviral fragment into the Colorado potato beetle or bracovirus genome (primer pairs 16-18).

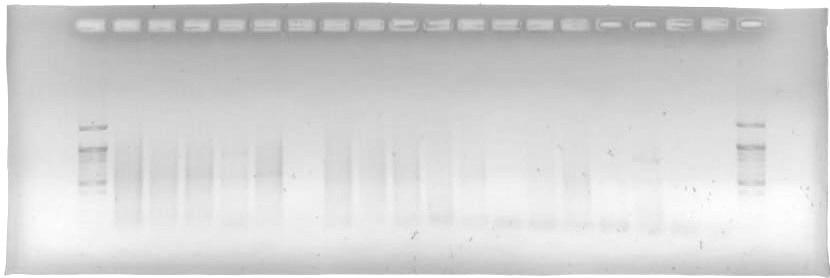

|  | **Br2Br_F1-Br2_ins_R1**  **(498 nt)** | | | | | | **Br2_ins_F1-Br2Br_R**  **(382 nt)** | | | | | | **Br2_ins_F1-Br2LD_R**  **(387 nt)** | | | | | |  |
| --- | --- | --- | --- | --- | --- | --- | --- | --- | --- | --- | --- | --- | --- | --- | --- | --- | --- | --- | --- |
| M | HL | E | PE | I | C | N | HL | E | PE | I | C | N | HL | E | PE | I | C | N | M |

M – marker

HL – hemolymph

E – eggs

PE – pure eggs

I – imago

C – cuticle

N – negative control
